# High-fidelity long-read metagenomics reveals the metabolic functions of dominant termite gut bacteria

**DOI:** 10.64898/2026.10.06.756956

**Authors:** Anna Prokhorova, Cong Liu, Vera Emelianenko, Soma Barawi, Simon Hellemans, Jan Šobotník, Gaku Tokuda, Yuichi Hongoh, Thomas Bourguignon

## Abstract

Many termites digest lignocellulosic wood using diverse gut microbiota, yet functions are rarely linked to specific symbionts because complete genomes are scarce. We used PacBio HiFi long reads to reconstruct circular single-contig bacterial metagenome-assembled genomes (MAGs) from the gut microbiota of the wood-feeding termite *Nasutitermes takasagoensis*. From 13.9 Gb of HiFi data, we obtained highly contiguous assemblies, including 11 *Breznakiellaceae* (*Spirochaetota*), two *Candidatus* Fibrimonas (*Fibrobacterota*), and one *Chitinispirillaceae* genome—three dominant lineages in this gut ecosystem. The reconstructed genomes encoded extensive carbohydrate-active enzyme repertoires and anaerobic fermentation pathways consistent with lignocellulose degradation. They also carried genes for essential amino acid biosynthesis, while nitrogen-fixation capacity was present in representatives of all three lineages but varied among closely related strains. One *Breznakiellaceae* genome possessed a complete Wood–Ljungdahl pathway potentially supporting reductive acetogenesis in the presence of external formate, whereas related strains carried incomplete variants likely involved in anabolic C□ metabolism. Differences in CAZymes, C□ metabolism, and nitrogen fixation among closely related strains reveal functional diversification that fragmented MAGs would not resolve. The mapping of metatranscriptomic reads from three biological replicates of the fiber-associated community to the reconstructed genomes provided independent evidence that key predicted metabolic functions are transcriptionally active. Our results demonstrate the power of HiFi metagenomics to recover highly contiguous genomes from complex insect microbiomes and uncover lineage-specific metabolic roles of dominant termite gut symbionts.

**IMPORTANCE:** Termites are among the most efficient degraders of plant biomass in nature and rely on specialized gut microbes to convert lignocellulose into nutrients. However, most termite gut bacteria remain uncultivated, limiting our understanding of their metabolic capabilities and ecological roles. Using PacBio HiFi long-read metagenomics, we reconstructed highly contiguous genomes of dominant termite gut bacteria, including the first complete genome from termite-associated *Candidatus* Fibrimonas. These genomes revealed complementary metabolic functions involved in lignocellulose degradation, reductive acetogenesis, nitrogen metabolism, and energy conservation. Metatranscriptomic data further showed that key predicted metabolic functions were transcriptionally active in the fiber-associated community. Our study shows how long-read metagenomics combined with transcriptomic evidence can resolve the metabolic roles and functional diversification of previously uncultivated microorganisms in complex microbial communities.

## INTRODUCTION

Termites are one of the most efficient decomposers of lignocellulose, the main component of wood (1). Lignocellulose is a complex polymer recalcitrant to enzymatic activities, consisting of cellulose, hemicellulose, lignin, and a variety of minor components (2). The decomposition of lignocellulose involves the action of a cocktail of carbohydrate-active enzymes (CAZymes), only a few of which are encoded in the genome of termites (3,4). Most lignocellulose-degrading CAZymes in the gut of termites are of microbial origin, produced by symbiotic bacteria, archaea, and protists (4). These gut microbes are ubiquitous in termites, except for protists that have been lost in the most diverse termite lineage, the Termitidae (1,5). Termite gut bacteria and archaea also contribute to the metabolism of their host in several additional ways. For example, they participate in nitrogen metabolism, allowing termites to thrive on wood, a substrate containing low amounts of nitrogen (6,7).

Our understanding of the termite gut microbiota and its function has been impeded by our inability to culture most termite gut bacteria (8). Many bacterial lineages are only known from DNA sequencing efforts (9). The function of these lineages is generally unknown. One approach to studying the taxonomy and function of uncultured bacteria is metagenomic analyses of entire gut microbial communities. Shotgun metagenomics has been performed on many termite species using short-read sequencing technology (10–12). However, because termites harbor hundreds of closely related bacterial lineages in their guts (13,14), short-read assemblies are often highly fragmented, limiting our abilities to assign functional capacities to specific bacteria. In addition, the fragmentation of genomes into multiple contigs can hinder the reconstruction of complete functional pathways, because their constituent genes may not be recovered within the same genome assembly. Metagenome-assembled genomes (MAGs), which group together metagenome contigs putatively part of an individual genome, partially address these problems. However, MAGs represent genomic bins that may encompass sequences from multiple closely related strains and often exhibit a certain degree of incompleteness and contamination (10,15). Complete genomes, particularly single-contig circular genomes, can overcome many of these limitations, enabling the inference of metabolic pathways from individual bacterial populations.

Most complete genomes of termite gut bacteria sequenced to date are those of protist endosymbionts obtained by sequencing of single protist cells (16–19). While useful, this approach does not allow to sequencing of bacteria living freely in the termite gut metagenomes assembled with short reads are **always** highly fragmented and cannot be applied to Termitidae, which host complex communities of bacteria composed of more than 1500 phylotypes (14). The reconstruction of complete bacterial genomes from termite guts may be possible using long-read sequencing technology, which has been successfully used with other complex bacterial communities (20,21). PacBio High Fidelity (HiFi) reads, which are over ten kilobase pairs and have error rates similar to those of short reads, appear especially promising for the reconstruction of highly contiguous metagenomes. HiFi reads allowed the reconstruction of complete bacterial genomes from the human gut microbiota (22), samples of sheep feces (23), and chicken cecum (24). The successful use of HiFi reads to reconstruct entire bacterial genomes from complex gut microbiota calls for their use to investigate other systems, such as the termite gut microbiota.

In this study, we used PacBio HiFi reads to sequence the whole gut microbiota of the wood-feeding termite *Nasutitermes takasagoensis* (Termitidae). HiFi read metagenome assemblies included bacterial genomes successfully reconstructed in one contig, which we used to determine the metabolic capabilities of the dominant gut bacterial lineages. These metabolic reconstructions allowed us to determine how the principal gut bacterial lineages of *N. takasagoensis* contribute to the digestion of lignocellulose and the nutrition of their host. They also revealed functional differences among closely related strains by comparing their metabolic capacities. We further integrated publicly available metatranscriptomic data from the fiber-associated bacterial community (25) to evaluate whether the metabolic functions inferred from the reconstructed genomes are transcriptionally active.

## RESULTS AND DISCUSSION

### HiFi reads allow the reconstruction of complete genomes of the dominant gut bacteria of *N. takasagoensis*

We obtained 1,090,744 PacBio HiFi reads from the whole gut of *N. takasagoensis*, making up a total of 13.9 Gb. HiFi reads, which included all reads with a quality threshold above Q20, had a median read quality value of Q32. We assembled HiFi reads with Flye and obtained 11,702 prokaryotic contigs longer than 20 kb, including 11,449 assigned to bacteria and 30 to archaea. The assembly included 50 contigs longer than 1 Mb and 12 circular bacterial genomes. We supplemented this Flye assembly with two additional bacterial genomes assembled with Hifiasm-meta. The 14 genomes included 11 genomes of the family *Breznakiellaceae* (phylum *Spirochaetota*), two genomes of *Candidatus (Ca.)* Fibrimonas (phylum *Fibrobacterota*), and one genome of the family *Chitinispirillaceae* (*Fibrobacterota*) (Table 1). All 14 genomes were recovered as circular single-contig assemblies. Thirteen had >90% estimated completeness, whereas *Ca.* Fibrimonas strain 2 had a lower estimated completeness of 79.85% despite being recovered as a circular single-contig assembly (Table 1). We retained both *Ca. Fibrimonas* strains for downstream analyses, while interpreting the absence of genes from strain 2 with caution (Table 1). We retained both *Ca.* Fibrimonas strains for downstream analyses, acknowledging that the absence of genes in strain 2 should be interpreted with caution. The completeness of 13 of our genomes was further supported by the long-read coverage. The sequencing depth of 10 kb sliding windows was consistent for all bacterial genomes (Figure S2A), suggesting negligible assembly errors. We computed the sequencing depth of 10 kb sliding windows once more using the original genome contigs broken at the midpoint with linked terminals. These analyses showed that the sequencing depth around the midpoint was consistent with other parts (Figure S2B), supporting the completeness of our genomes. The exclusive recovery of *Spirochaetota* and *Fibrobacterota* genomes reflects the high relative abundance of these two phyla in *Nasutitermes* (10,26,27). In our dataset, *Spirochaetota* and *Fibrobacterota* were represented by 55.5% and 11.3% of the prokaryotic reads, respectively (Figure 1A). In contrast, other bacterial phyla were represented by less than 5% of the prokaryotic reads and were not assembled into complete genomes because of insufficient coverage (Figure 1B). For comparison, two previously generated Illumina termite-gut metagenomes (BDIT094 and THAI067, (10)) yielded 5 and 2 MAGs with ≥50% completeness and <10% contamination from 23.15 and 9.08 Gb of sequence data, respectively, corresponding to approximately 2.2 qualifying MAGs per 10 Gb in both datasets. In contrast, the 13.9-Gb HiFi dataset yielded 22 MAGs meeting the same quality threshold (15.8 per 10 Gb), 18 of which (81.8%) were single-contig assemblies, whereas none of the qualifying short-read MAGs were single-contig. Overall, our results demonstrated that HiFi reads allow the reconstruction of highly contiguous genomes of the dominant microbial members of a complex termite gut microbiota. Collectively, the 14 reconstructed genomes accounted for 60.1% of the HiFi reads at MAPQ ≥20, with individual genomes accounting for 1.82–9.64% of the total reads (Table 1).

**Figure 1.**
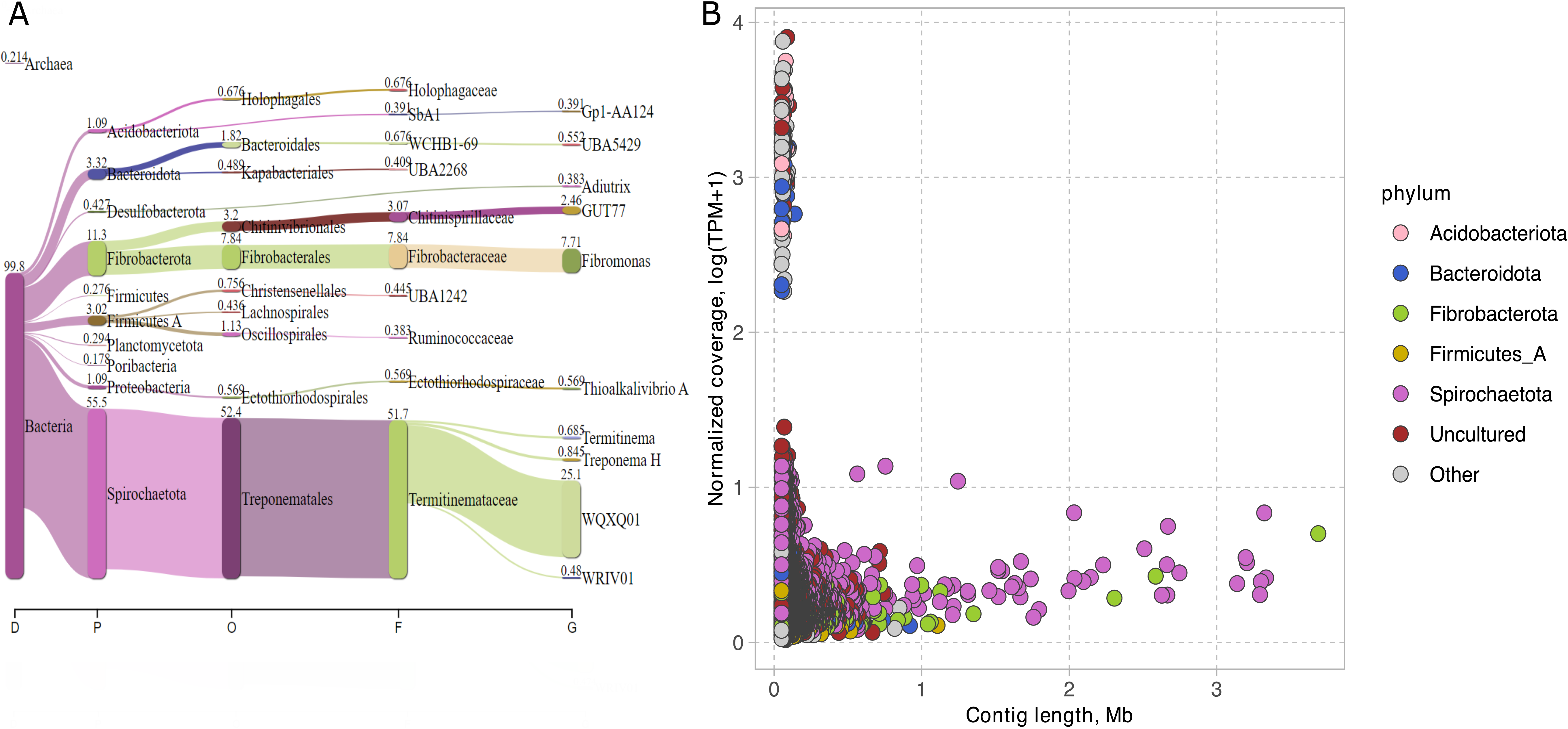
Taxonomical composition of the *N. takasagoensis* gut microbiome. A) Sankey diagram depicting the distribution of the bacterial taxa in the gut of *Nasutitermes takasagoensis*. Species are ranked by the Genome Taxonomy Database (GTDB); B) Distribution of contig lengths (X-axis) and their coverage depth (Y-axis) in the metagenome assembly of *Nasutitermes takasagoensis* gut microbiome.

**Table 1.** Genomic characteristics and representation of the 14 circular bacterial genomes reconstructed from the *N. takasagoensis* gut metagenome. All genomes were recovered as circular single-contig assemblies. Completeness and contamination were estimated using CheckM2. Mapped HiFi reads (%) represents the percentage of the HiFi reads mapped competitively to each of the 14 reconstructed genomes, retaining primary alignments with MAPQ ≥20.

| GENOMES | Completeness [%] | Contamination [%] | GC [%] | Coding density [%] | rRNA | tRNA | CDCs | Mapped HiFi reads [%] | Genome size [Mbp] |
| --- | --- | --- | --- | --- | --- | --- | --- | --- | --- |
| *Breznakiellaceae* strain 1 | 100 | 0.71 | 46 | 88.59 | 6 | 47 | 4217 | 4.04 | 4.24 |
| *Breznakiellaceae* strain 2 | 99.4 | 2.44 | 49 | 93.5 | 8 | 45 | 3169 | 3.64 | 3.33 |
| *Breznakiellaceae* strain 3 | 94.15 | 1.4 | 48 | 91.92 | 6 | 46 | 2540 | 2.1 | 2.61 |
| *Breznakiellaceae* strain 4 | 99.3 | 0.9 | 43 | 92.58 | 2 | 49 | 3180 | 2.88 | 2.65 |
| *Breznakiellaceae* strain 5 | 100 | 0.86 | 40 | 90.85 | 3 | 46 | 2522 | 2.69 | 3.25 |
| *Breznakiellaceae* strain 6 | 99.26 | 4.8 | 41 | 90.21 | 2 | 48 | 2478 | 3.06 | 3.23 |
| *Breznakiellaceae* strain 7 | 96.55 | 3.69 | 43 | 92.3 | 6 | 47 | 3110 | 3.27 | 2.79 |
| *Breznakiellaceae* strain 8 | 96.31 | 0.09 | 45 | 90.64 | 6 | 47 | 2593 | 3.74 | 3.27 |
| *Breznakiellaceae* strain 9 | 98.3 | 1.91 | 45 | 90.82 | 6 | 46 | 3035 | 3.15 | 3.2 |
| *Breznakiellaceae* strain 10 | 97.33 | 2.07 | 44 | 90.78 | 6 | 47 | 2940 | 6.03 | 3.19 |
| *Breznakiellaceae* strain 11 | 100 | 0.62 | 45 | 91.01 | 6 | 46 | 2897 | 6.21 | 2.67 |
| *Co. Fibrimonas* strain 1 | 97.6 | 0.58 | 44 | 94.5 | 4 | 45 | 2558 | 2.3 | 2.51 |
| *Co. Fibrimonas* strain 2 | 79.85 | 6.86 | 43 | 94.12 | 2 | 35 | 2403 | 1.82 | 2.3 |
| *Chitinophagaceae* strain 1 | 99.01 | 0.4 | 54 | 96.93 | 6 | 46 | 3232 | 3.46 | 3.65 |

### Phylogenetic position of 14 newly reconstructed bacterial genomes

We reconstructed 14 circular single-contig bacterial genomes, comprising 11 Spirochaetota and three Fibrobacterota genomes (Table 1). Thirteen of the 14 genomes had >90% estimated completeness according to CheckM2, whereas *Ca. Fibrimonas* strain 2 had a lower estimated completeness of 79.85% despite being recovered as a circular single-contig assembly. To place these genomes into an evolutionary context, we reconstructed a maximum-likelihood phylogenetic tree alongside representative genomes and metagenome-assembled genomes (MAGs) from termite guts available in GTDB, including MAGs reported by Hervé et al. (2020) and Mies et al. (2024) (Figure 2). Our 11 genomes of Spirochaetota formed several lineages within the family *Breznakiellaceae*, a termite-associated clade previously shown to co-speciate with termites over evolutionary timescales (28–30). This termite-specific clade is phylogenetically related to *Breznakiella*, a genus of Spirochaetota isolated from the Madeira cockroach *Rhyparobia maderae* (31), indicating its long history of association with insect guts. Most strains clustered within the lineage *WQXQ01* together with MAGs by Hervé et al. (2020) and Mies et al. (2024), whereas strain 1 formed a separate lineage more closely related to *Leadbettera azotonutricia*, which was isolated from the gut of the termite *Zootermopsis angusticollis* (32). Relative evolutionary divergence (RED) values and phylogenetic distances supported the separation of *WQXQ01* from *Leadbettera* sensu stricto.

**Figure 2.**
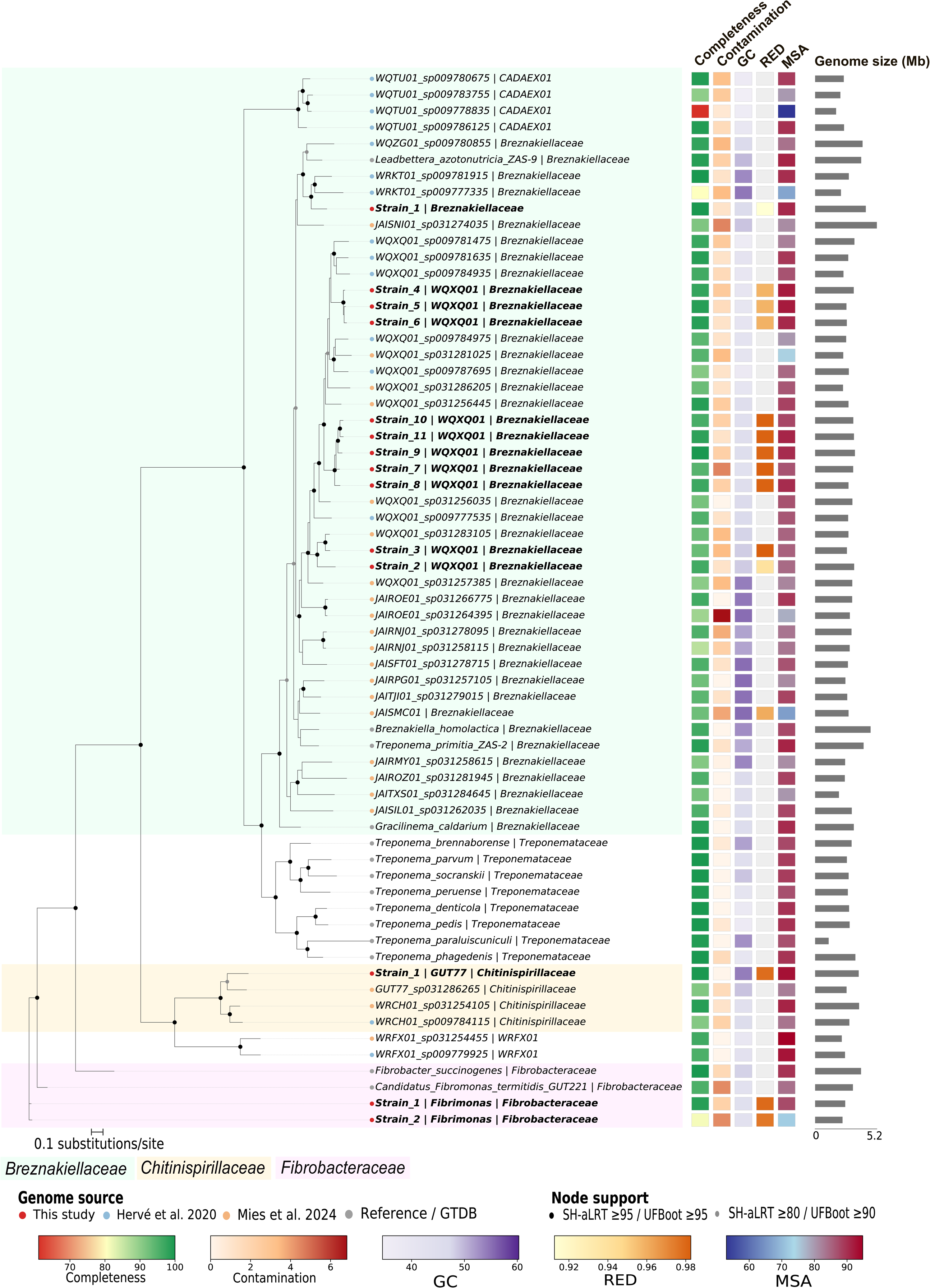
Phylogenomic placement of genomes reconstructed from the gut microbiome of *Nasutitermes takasagoensis*. Maximum-likelihood phylogeny of representative members of the families *Breznakiellaceae, Chitinispirillaceae*, and *Fibrobacteraceae* based on conserved marker proteins. Genomes reconstructed in this study are highlighted in red. Background shading indicates family-level assignments. Filled circles indicate branch support (SH-aLRT/UFBoot). The accompanying heatmap shows genome completeness, contamination, GC content, relative evolutionary divergence (RED), and marker set alignment (MSA) values, while gray bars indicate genome size. Scale bar represents 0.1 substitutions per site.

Average Nucleotide Identity (ANI) analyses further supported substantial genomic diversification within *Breznakiellaceae*. Pairwise ANI values among the *Breznakiellaceae* genomes reconstructed in this study ranged from 79.0% to 91.1% (Table S1), indicating that the strains represent multiple distinct species. ANI values between *WQXQ01* strains and *Leadbettera azotonutricia* were below genus-level thresholds proposed for prokaryotic taxonomy, consistent with their placement into separate genus-level lineages. These results highlight the extensive diversification of termite-associated *Breznakiellaceae* and suggest that closely related lineages occupying the same gut environment may possess distinct metabolic capacities.

The Fibrobacterota genomes reconstructed in this study belonged to two termite-specific clades known to coevolve with their hosts (28), for which previously available genomes were represented by fragmented MAGs. The two *Ca.* Fibrimonas clustered within the termite-associated *Fibrobacteraceae* lineage that includes *Ca.* Fibrimonas termitidis GUT221 described by Abdul Rahman et al. (33), an uncultured genus endemic to termite gut and primarily found in Termitidae (28,33), having for sister group *Fibrobacter succinogenes*, a bacterial species isolated from the rumen of cattle (34). ANI values between our two *Ca.* Fibrimonas genomes were approximately 91.7%, supporting their classification as separate congeneric species. Whereas ANI values relative to *Ca.* Fibrimonas termitidis were ∼77%, indicating substantial divergence.

The last genome belonged to a clade of *Chitinispirillaceae* related lineage GUT77 endemic to the gut of Termitidae (10,28) and distantly related to *Chitinivibrio alkaliphilus*, a bacterium isolated from hypersaline lake sediments (35). Although some genomes from these lineages were recovered as MAGs using Illumina short-read sequencing by Abdul Rahman et al. (33), they remained fragmented and incomplete. The circular single-contig genomes reconstructed here substantially improve genome contiguity for these termite-specific Fibrobacterota lineages compared with previously available fragmented MAGs.

### Transcriptional activity of reconstructed genomes in the fiber-associated community

To evaluate whether the reconstructed genomes represent metabolically active members of the *N. takasagoensis* gut microbiota, we analyzed metatranscriptomic data from three previously published biological replicates of the fiber-associated bacterial community in the termite hindgut (BioProject PRJDB5939) (25). Transcriptomic reads were mapped onto the 14 reconstructed genomes, yielding overall alignment rates of 46.5–47.5% across the three biological replicates (Table S5). Approximately 11 million paired-end reads were assigned to protein-coding genes in each replicate, indicating that the reconstructed genomes represent a substantial fraction of the transcriptionally active fiber-associated bacterial community.

Gene expression profiles were congruent among biological replicates. Transcripts were detected from all 14 reconstructed genomes, although expression levels varied among lineages and metabolic pathways (Figure S1). Genes involved in carbohydrate degradation, hydrogen metabolism, WLP/C1 metabolism, nitrogen fixation, and ammonium assimilation were transcriptionally active across the genomes encoding these functions. Pathway-specific expression patterns are discussed below in the context of the corresponding metabolic reconstructions. Overall, the metatranscriptomic data provide independent evidence that the reconstructed genomes represent metabolically active members of the fiber-associated community and support the physiological relevance of several metabolic functions inferred from their genomic repertoires.

### Genes encoding enzymes potentially involved in lignin modification/degradation or oxygen tolerance in the gut microbiota of *N. takasagoensis*

We evaluated the metabolic potential of our reconstructed circular bacterial genomes using Prokka and KEGG annotations, with a particular focus on the degradation of lignin, the most recalcitrant component of lignocellulose. Although lignin depolymerization in the termite gut remains poorly understood, previous studies have reported signs of microbial involvement in this process (36,37). We therefore searched our 14 genomes for lignin-modifying enzymes (LME) and lignin-degrading auxiliary (LDA) enzymes, such as laccases, manganese peroxidases, and cellobiose dehydrogenases, which are commonly associated with lignin breakdown in aerobic or facultatively anaerobic organisms.

We identified several genes in our bacterial genomes encoding putative LMEs and LDAs. For example, *Breznakiellaceae* strains 1 and 5 encoded manganese-dependent peroxidase (EC:1.11.1.13), and strains 3, 10, and 11 encoded laccase (EC:1.10.3.2). Pyranose 2-oxidases (EC:1.1.3.10) were present in most *Breznakiellaceae* genomes, both strains of *Ca.* Fibrimonas, and *Chitinispirillaceae* strain 1. Cellobiose dehydrogenase (EC:1.1.99.18) was found in all 14 genomes (Figure 3 and Table S2). *Breznakiellaceae* strain 1 also encoded vanillyl-alcohol oxidase (EC:1.1.3.38, CAZyme AA4 family). Additional LMEs and LDAs, such as lignin peroxidase (EC:1.11.1.14), dye-decolorizing peroxidase (EC:1.11.1.19), and glyoxal oxidase (EC:1.2.3.15) were detected in genome fragments assigned to *Candidatus* Adiutrix, *Ca.* Fibrimonas and other *Fibrobacterales*, *Treponematales*, and some uncultured taxa. However, even though these genes are homologous to known lignin-degrading enzymes, their presence does not provide irrefutable proof of lignin-degrading activity.

**Figure 3.**
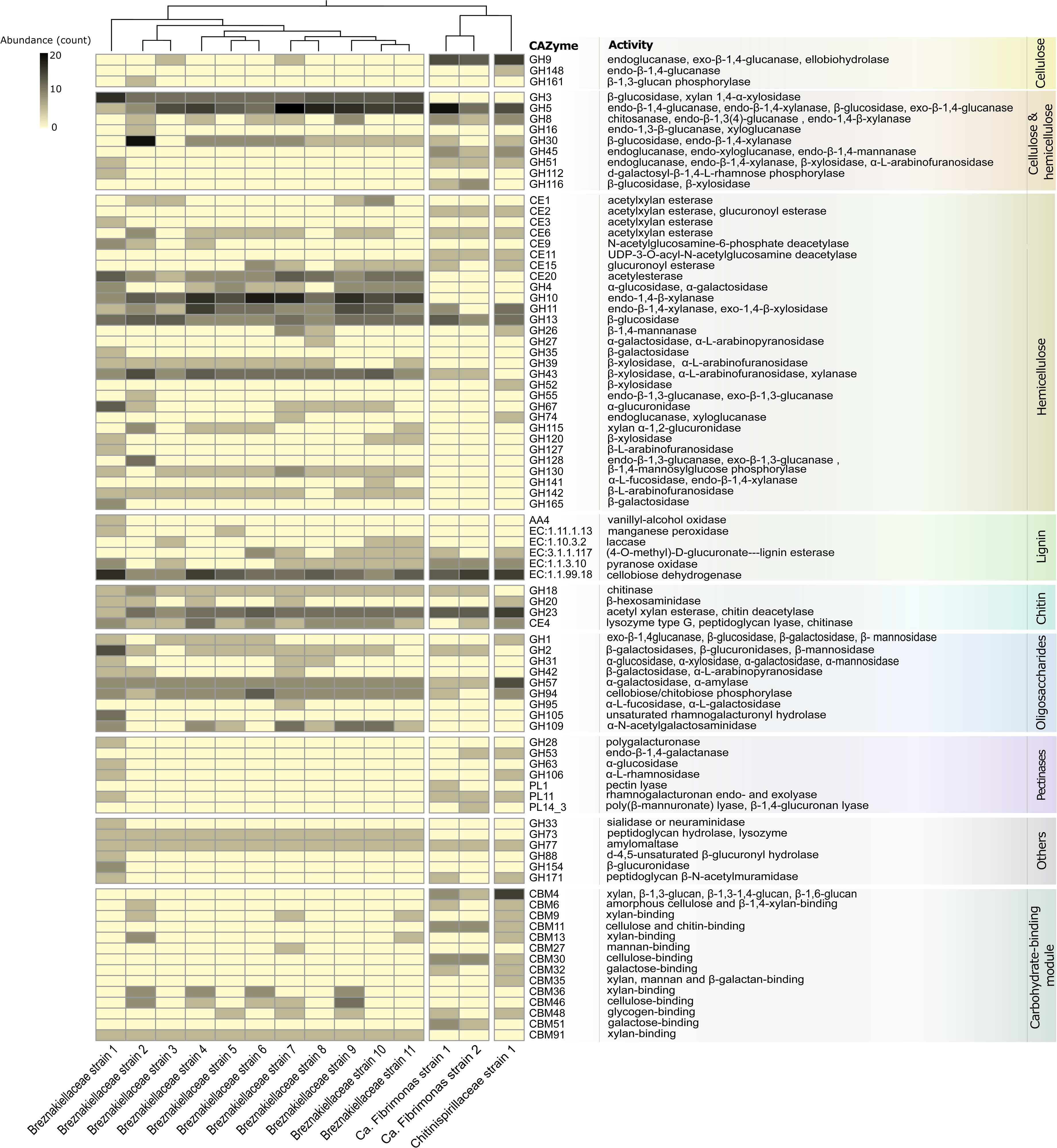
Distribution of carbohydrate-active enzymes (CAZymes) in genomes reconstructed from the gut microbiome of *Nasutitermes takasagoensis*. Heatmap showing the abundance of glycoside hydrolases (GHs), carbohydrate esterases (CEs), auxiliary activities (AAs), polysaccharide lyases (PLs), and carbohydrate-binding modules (CBMs) identified in *Breznakiellaceae*, *Ca.* Fibrimonas, and *Chitinispirillaceae* genomes. CAZyme families are grouped according to their predicted substrate specificity and biological function, including cellulose, hemicellulose, lignin, chitin, oligosaccharide, and pectin degradation, as well as carbohydrate-binding activities. Color intensity indicates the number of genes assigned to each CAZyme family. Hierarchical clustering was performed based on CAZyme abundance profiles.

These enzymes require oxygen or hydrogen peroxide to function and need to be excreted extracellularly, necessitating secretion signals and the appropriate redox context. *L. azotonutricia* and *T. primitia*, the closest cultured relatives of our *Breznakiellaceae* genomes, as well as the *Ca.* Fibrimonas MAGs reported by Abdul Rahman et al. (33), lack classical oxidative stress enzymes such as catalase or superoxide dismutase. In our dataset, neither *Ca.* Fibrimonas 1 nor *Ca.* Fibrimonas 2 encoded these enzymes, but both possessed cytochrome bd-type ubiquinol oxidase, which can remove oxygen under microoxic conditions, as well as glutathione peroxidase (EC:1.11.1.9). Many genomes, including the *Ca.* Fibrimonas strains, also contained homologs annotated as L-threonate 2-dehydrogenase (EC:1.1.1.411), which may correspond to NAD(P)H oxidase or represent related enzymes. Therefore, like in *T. primitia*, in which NAD(P)H oxidase and NAD(P)H peroxidase act as alternative antioxidant systems (38), LMEs and LDA may primarily serve as antioxidants rather than lignin degradation. Our results do not entirely rule out the possibility that some gut bacteria contribute to lignin modification to facilitate access to cellulose and hemicellulose; however, additional evidence is needed to support this hypothesis.

### *Breznakiellaceae, Candidatus* Fibrimonas, and *Chitinispirillaceae* degrade cellulose and hemicellulose in synergy

The degradation of cellulose and hemicellulose normally requires the synergistic action of a broad array of CAZymes. We identified CAZymes across our 14 genomes using dbCAN (39) and the CAZy database (40). Our 14 genomes included 100 CAZyme families, including 55 families of glycosyl hydrolases (GHs), 18 families of glycoside transferases (GTs), 14 families of carbohydrate binding modules (CBMs), nine families of carbohydrate esterases (CEs), three families of polysaccharide lyases (PLs), and one family of auxiliary activity (AA) (Figure 3). We primarily focused our analyses on CAZymes involved in cellulose and hemicellulose degradation (GHs, CEs, PLs, and AAs) and binding to specific carbohydrates (CBMs).

The complete hydrolysis of cellulose, a linear chain of glucose, requires the cooperation of endoglucanases (EC 3.2.1.4), exoglucanases (EC 3.2.1.91 and EC 3.2.1.176), and β-glucosidases (EC 3.2.1.21) (41). Endoglucanase genes were unequally distributed, as they were primarily found in the genomes of *Ca*. Fibrimonas and *Chitinispirillaceae*. Our three genomes of *Fibrobacterota* contained between four and six copies of GH9, a family of cellulases with endoglucanase activity known to synergistically increase the hydrolytic ability of other cellulases (42). The genomes of *Ca.* Fibrimonas and *Chitinispirillaceae* also contained at least one copy of GH8, GH45, and GH51, three enzymes with endoglucanase activities (42). Consistent with these predictions, genes involved in cellulose degradation were actively transcribed in all three metatranscriptomes (Figure S1, Table S6). However, we found no decisive evidence for the presence of exoglucanases, which act on the terminals of oligosaccharide chains generated by endoglucanases, releasing glucose or cellobiose. Each genome of *Ca*. Fibrimonas and *Chitinispirillaceae* contained between seven and thirteen GH5-family enzymes/genes, a CAZyme family known to sometimes, but not always, have exoglucanase activities (43).

Therefore, it is unclear whether *Ca.* Fibrimonas and *Chitinispirillaceae* use exoglucanase to degrade cellulose, but their absence could be compensated by a high number of β-glucosidases and endoglucanases, as Allardyce et al. (44) proposed in their model of cellulose hydrolysis in the land crab *Gecarcoidea natalis*. In support of this hypothesis, the main cellulases found in the genome of *Fibrobacter succinogenes,* which is capable of growth on crystalline cellulose (45), are GH8, GH9, GH45, and GH51 (42), four CAZymes present in all three genomes of *Ca.* Fibrimonas and *Chitinispirillaceae*.

All our 14 genomes contained one to 13 copies of GH5, one of the most diverse CAZyme families, which is divided into more than 50 subfamilies with various catalytic activities (43). The most abundant GH5 subfamily across *Breznakiellaceae* strains was GH5_4, which has high catalytic activity for hemicellulose (46). In contrast, the GH5 subfamily primarily found in *Ca.* Fibrimonas and *Chitinispirillaceae* was GH5_2, a subfamily with cellulolytic activity (Table S2). In addition, CBM4, CBM6, and CBM11, which are usually associated with the GH5 family and bind cellulose (43), were highly abundant in *Ca*. Fibrimonas and *Chitinispirillaceae* genomes. Therefore, the GH5 repertoire of *Breznakiellaceae* was consistent with a specialization on hemicellulose, while that of *Ca.* Fibrimonas and *Chitinispirillaceae* indicated a specialization on cellulose.

Although the analysis of the CAZyme repertoires of our 14 genomes indicates that cellulose degradation is primarily performed by *Ca.* Fibrimonas and *Chitinispirillaceae*, several strains of *Breznakiellaceae* contained a few glucanases in addition to β-glucosidases and GH94 phosphorylases, suggesting they participate, to a certain extent, in cellulose degradation. For example, the genome of *Breznakiellaceae* strain 2 contained one GH8 and one endoglucanase from the GH16 family that was not found in the other 13 genomes. The genome of *Breznakiellaceae* strain 7 contained two endoglucanases from GH8 and GH9 families. Other genomes of *Breznakiellaceae* contained a maximum of three endoglucanases. These results indicate that *Breznakiellaceae*, which is known to primarily specialize in hemicellulose (25), has the potential to break down cellulose, albeit not to the extent of *Ca.* Fibrimonas and *Chitinispirillaceae*.

In contrast to endoglucanases, β-glucosidases were found in most of our 14 genomes. However, while all *Breznakiellaceae* genomes possessed multiple GH3 β-glucosidases, this enzyme family was absent in *Ca*. Fibrimonas and *Chitinispirillaceae*. Instead, *Ca.* Fibrimonas genomes contained GH116 β-glucosidases, while the genome of *Chitinispirillaceae* showed no apparent presence of β-glucosidases (Figure 3). *Chitinispirillaceae* and *Ca.* Fibrimonas strain 1 encoded intracellular GH94 phosphorylases, an enzyme that cleaves cello-oligosaccharides to produce glucose-1-phosphates. Therefore, we postulate that these strains directly adhere to wood fibers, which they hydrolyze, producing cello-oligosaccharides that they transport in the cytoplasm, as is the case in many anaerobic cellulolytic bacteria (2). Overall, the high abundance of cellulases in the CAZyme repertoire of *Ca.* Fibrimonas and *Chitinispirillaceae* supports cellulose as their primary source of carbon.

The hydrolysis of hemicellulose, a polysaccharide composed of diverse sugars linked by complex networks of bonds, requires the action of various debranching enzymes. While our 14 genomes all contained hemicellulases, these enzymes were particularly abundant and diverse in the genomes of *Breznakiellaceae* (Figure 3). In total, *Breznakiellaceae* genomes contained between 17 and 40 CAZymes specifically degrading hemicellulose, while *Ca*. Fibrimonas and *Chitinispirillaceae* contained between 9 and 23 such CAZymes (Table S2).

The main polysaccharides present in wood hemicellulose are glucurono(arabino)xylans and (galacto)glucomannans, which have backbones of xylans and glucomannans, respectively (47). The genomes of *Breznakiellaceae* contained genes encoding enzymes that degrade both xylans and glucomannans. The most abundant hemicellulases were xylanases, which hydrolyze xylan, and were represented by multiple CAZyme families, including four found across all genomes of *Breznakiellaceae*: GH3, GH10, GH11, and GH43. Multiple other xylanases, including potential xylosidases such as GH39, were also found in a subset of *Breznakiellaceae* genomes. Additionally, each *Breznakiellaceae* genome contained accessory or debranching enzymes needed for the complete degradation of complex substituted xylans, such as glucuronoarabinoxylans. These CAZymes included α-fucosidase (GH3, GH95, and GH141) and arabinofuranosidase (GH2, GH43, GH51, GH127, and GH142). Glucomannans are hydrolyzed by a combination of endo-β-1,4-mannanase and endo-β-1,4-glucanases. Most *Breznakiellaceae* strains possessed several GH5_4, including endo-β-1,4-mannanases and glucomannan-specific endo-β-1,4-glucanase. Therefore, the hemicellulase repertoire of *Breznakiellaceae* strains suggests they are capable of degrading the two major hemicellulosic backbones present in the wood. With some exceptions, such as GH11 and GH43, most hemicellulases were not encoded in the genomes of *Ca.* Fibrimonas and *Chitinispirillaceae*, which aligns with their specialization in cellulose degradation. Overall, our results are consistent with a previous study that, based on transcriptomic data, identified *Breznakiellaceae* as the major contributor to hemicellulose degradation in the gut of *N. takasagoensis* (25). However, by using complete genomes rather than transcriptomes, we could assess pathway completeness, CAZyme subfamily composition, and the genomic context, revealing additional strain-level differences and gene absences that are difficult to infer from transcriptomes alone. In the gut of *N. takasagoensis*, the monosaccharides composing lignocellulose are primarily released by the synergistic action of the cellulolytic *Ca.* Fibrimonas and *Chitinispirillaceae* and the hemicellulolytic *Breznakiellaceae*. Additionally, the presence of genes involved in the cleavage of uronic acid bonds (GH4, GH28, GH67, GH88, GH105, GH106, GH115, PL1, PL11, and PL14) indicates a genomic potential to degrade pectin, a major component of primary plant cell walls.

### Breznakiellaceae, Candidatus Fibrimonas, and Chitinispirillaceae ferment wood polysaccharides

The hindgut of termites is microoxic at its periphery and anoxic in the center, where the microbial fermentation of wood polysaccharides generates acetate, lactate, ethanol, CO_2_, and H_2_ (1,8). Termites primarily use acetate as a source of energy (48). We examined the 14 reconstructed bacterial genomes for pathways involved in fermentation and anaerobic respiration (Figures 4 and 5). All genomes encoded some components of the central carbon metabolism, including an often-partial Embden–Meyerhof pathway (glycolysis), the pentose phosphate pathway, and an incomplete tricarboxylic acid (TCA) cycle (Figure 4, Table S3). Notably, 2-oxoglutarate dehydrogenase were absent from all genomes (Table S3), which is consistent with the truncation of the oxidative branch of the TCA cycle in many anaerobes (49). Genes for the Entner–Doudoroff pathway (*edd* (EC:4.2.1.12)*, eda* (EC:4.2.1.14), and *kdgK* (EC:2.7.1.45)) were found in most genomes, but none of the *Breznakiellaceae* strains encoded glucose-6-phosphate 1-dehydrogenase (Zwf). Therefore, the Entner–Doudoroff pathway is incomplete in all *Breznakiellaceae*, and the likely function of these genes is uronate degradation. *Chitinispirillaceae* strain 1 lacked enolase (Eno), which blocks the lower glycolytic pathway via the Embden–Meyerhof route. The mechanism used to bypass this step remains unknown, as discussed for other strict anaerobes (50).

**Figure 4.**
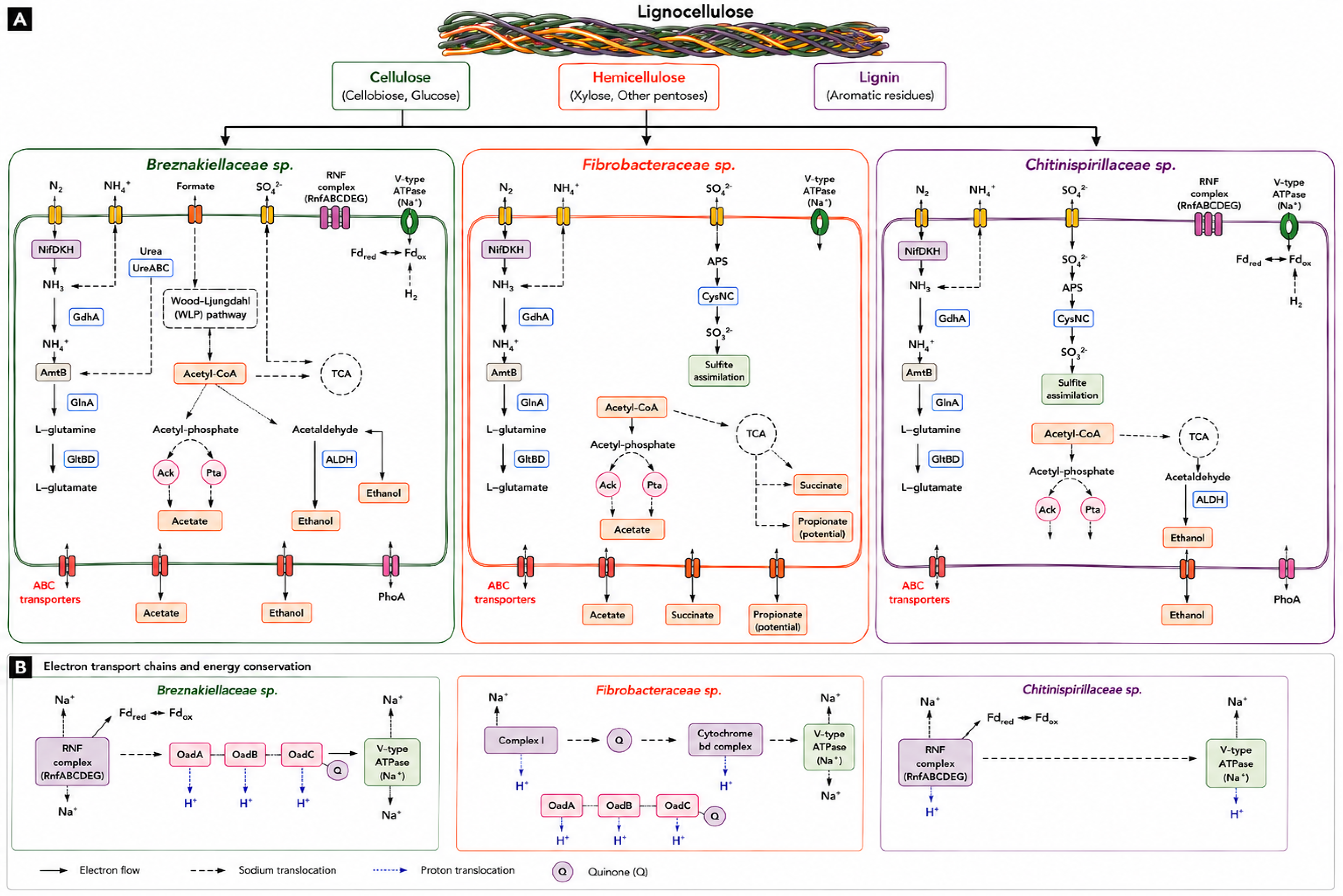
Predicted metabolic functions and energy conservation strategies of dominant termite gut bacteria reconstructed from the gut microbiome of the wood-feeding termite *Nasutitermes takasagoensis*. **(A)** Genome-based metabolic reconstruction of three dominant bacterial lineages, represented by *Breznakiellaceae* sp., *Fibrobacteraceae* sp., and *Chitinispirillaceae* sp. The model highlights complementary roles in lignocellulose utilization, nitrogen acquisition and recycling, sulfur metabolism, reductive acetogenesis, fermentation, and nutrient exchange within the termite gut ecosystem. **(B)** Predicted electron transport chains and energy conservation mechanisms encoded by the reconstructed genomes. *Breznakiellaceae* sp. and *Chitinispirillaceae* sp. possess RNF complexes coupled to V-type ATPases, suggesting sodium- and proton-dependent ion translocation linked to energy conservation. *Fibrobacteraceae* sp. encodes a respiratory chain comprising Complex I, quinone-dependent electron transfer, and a cytochrome *bd* oxidase complex, together with V-type ATPase and membrane-bound oxidoreductases (OadABC). Solid arrows indicate predicted electron flow, dashed arrows indicate sodium translocation, and blue dotted arrows indicate proton translocation.

**Figure 5.**
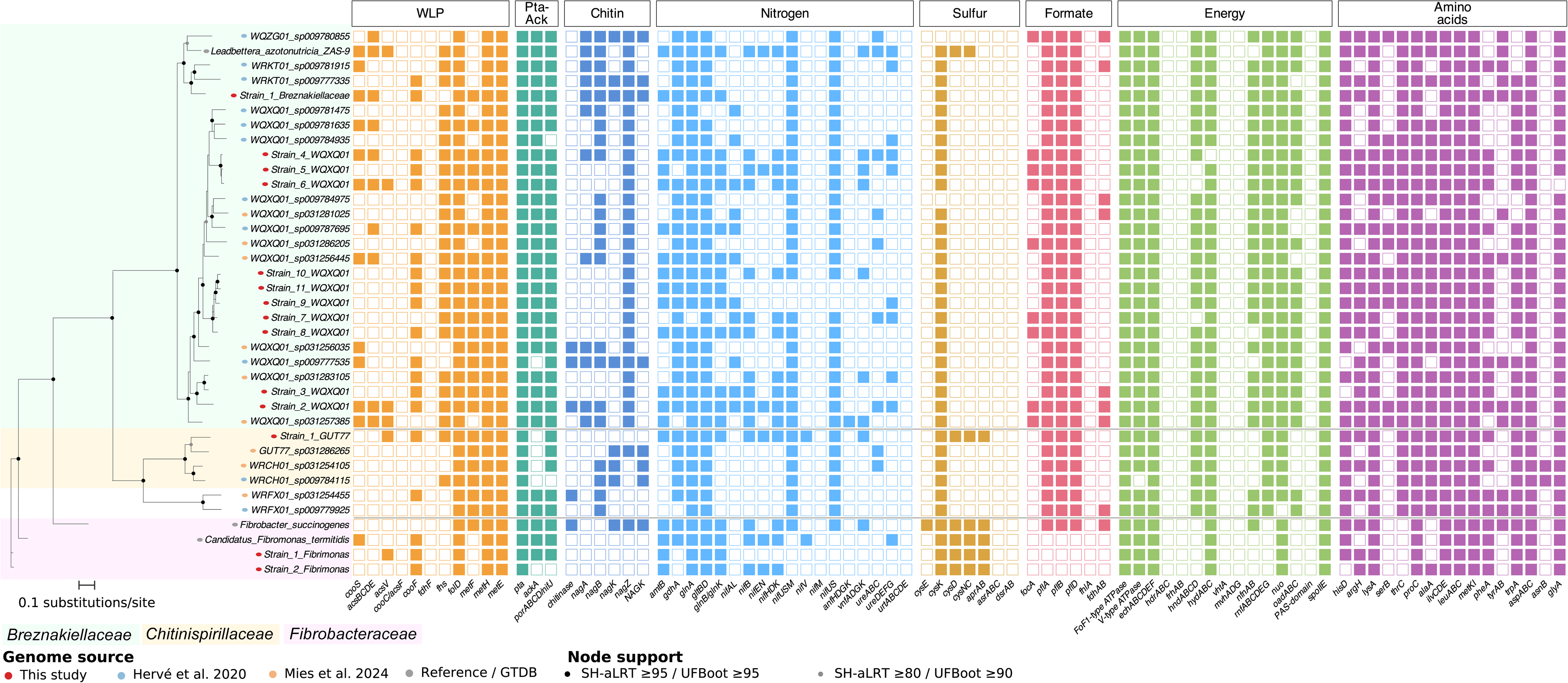
Distribution of metabolic traits among reconstructed termite gut genomes. Phylogenomic tree and presence–absence matrix of genes involved in the Wood–Ljungdahl pathway (WLP), acetate production (Pta–Ack pathway), chitin degradation, nitrogen metabolism, sulfur metabolism, formate metabolism, energy conservation, and amino acid biosynthesis in representative members of the families *Breznakiellaceae*, *Chitinispirillaceae*, and *Fibrobacteraceae*. Genomes reconstructed in this study are highlighted in red. Colored squares indicate gene presence, whereas empty squares indicate absence. Background shading denotes family-level taxonomic assignments. Branch support values are indicated by filled circles.

None of the *Breznakiellaceae*, *Candidatus* Fibrimonas, and *Chitinispirillaceae* genomes encoded L-lactate dehydrogenase [EC:1.1.1.27] or lactate-utilization enzymes, and no genes coding for pyruvate formate-lyase or formate dehydrogenase were detected. In *Ca.* Fibrimonas strain 1, sugars are fermented to acetate and succinate, and potentially to propanoate. NADH can be re-oxidized through complex I, with electrons transferred via the quinone pool to fumarate reductase (yielding succinate) or cytochrome bd-type ubiquinol oxidase (yielding H□O). The Na-transporting oxaloacetate/methylmalonyl-CoA decarboxylase OadABC [EC:7.2.4.2] can decarboxylate methylmalonyl-CoA, and the presence of an adjacent methylmalonyl-CoA epimerase gene [EC:5.1.99.1] in the genome suggests that succinyl-CoA, produced via the partial TCA cycle that includes succinate dehydrogenase/fumarate reductase SdhABC, is further metabolized to propanoate. OadABC exports Na, while complex I exports H□, together establishing a dual ion-motive force that drives ATP synthesis via a (Na□/H□-dependent) V-type ATPase. *Ca.* Fibrimonas strain 2 appears to encode the same pathway but lacked some genes, possibly due to genome incompleteness. Both *Ca.* Fibrimonas strains also encode cytochrome bd-type ubiquinol oxidase, which may facilitate oxygen removal under microoxic conditions.

*Chitinispirillaceae*, which lacked both complex I and OadABC, instead encoded a Na□-translocating ferredoxin:NAD oxidoreductase RNF complex (RnfABCDGE), likely generating a Na□ gradient for ATP production via a Na□-dependent V-type ATPase. It also possessed an NAD(P)-reducing iron hydrogenase HndABCD, probably cytoplasmic, which may oxidize NAD(P)H and ferredoxin to produce H_2_, and could be involved in signal transduction. In addition, *Chitinispirillaceae* encoded a putative sensory [FeFe]-hydrogenase, whose operon comprised a NuoE-like subunit, an iron hydrogenase B1/B3 catalytic domain-containing protein, and a sensory iron hydrogenase PAS-domain-containing catalytic subunit. In *Breznakiellaceae*, the main hydrogenase was the cytoplasmic, electron-bifurcating [FeFe]-hydrogenase HydABC, present in all strains except *Breznakiellaceae* strain 4, which instead possessed HndABCD. Many *Breznakiellaceae* strains (strains 1–3 and 8–11) also encoded a sensor-type HndABCD with an ATP-binding protein, while others (strains 4–7) possess an alternative sensory [FeFe]-hydrogenase complex with a PAS-domain-containing catalytic subunit and associated regulatory proteins. Hydrogen metabolism genes were among the most highly expressed pathways in all reconstructed lineages (Figure S1), supporting their central role in energy conservation.

Alcohol fermentation genes, such as those encoding aldehyde dehydrogenase and alcohol dehydrogenase, were present across all genomes of *Breznakiellaceae* and *Chitinispirillaceae*, whereas *Ca.* Fibrimonas did not possess these genes. All *Breznakiellaceae* strains encoded pathways for the fermentation of sugars to H□, CO□, acetate, and ethanol. Most strains use HydABC for hydrogen metabolism, while others use HndABCD, which includes an ATP-binding protein and is also found in *L. azotonutricia* (51). Some strains additionally encode sensory [FeFe]-hydrogenases that may function in H□ signaling. Acetate kinase (Ack), which catalyzes the interconversion between acetylphosphate and acetate, was present in all *Breznakiellaceae* genomes and in *Ca.* Fibrimonas strain 1.

Our three *Fibrobacterota* genomes did not encode true dissimilatory sulfite reductase (DsrAB) but instead contained genes consistent with assimilatory sulfite reduction, indicating their role in incorporating sulfite for biosynthesis rather than energy conservation. All *Breznakiellaceae* strains possessed the RNF complex for Na□/H□ motive force generation, and most also encoded OadABC for Na□.

The presence of putative cellulobiose phosphorylase (EC:2.4.1.20) and glucokinase (EC:2.7.1.2) across all genomes suggests the utilization of cellobiose, a major breakdown product of cellulose (Figure 4). This supports the role of *Breznakiellaceae, Candidatus* Fibrimonas, and *Chitinispirillaceae* as anaerobic fermenters of lignocellulosic carbohydrates with distinct variations in central metabolism reflecting their adaptation to the anoxic conditions prevalent in the gut of *N. takasagoensis*.

### *Breznakiellaceae* encode a non-canonical reductive acetogenesis pathway

Reductive acetogenesis, which converts H□ and CO□ into acetate via the Wood–Ljungdahl pathway (WLP), is considered a key hydrogen sink in wood-feeding termite guts (52–54). Although *Spirochaetota* are often seen as the principal acetogens in termite guts, this function has only been demonstrated in two species of *Spirochaetota*: *T. primitia* (32) and *Candidatus* Treponema intracellularis, an endosymbiont of the *Hodotermopsis sjoestedti* gut protist *Eucomonympha* (55). The 11 genomes of *Breznakiellaceae* encoded several components of the WLP, but only strain 2 possessed a complete pathway potentially capable of energy-conserving reductive acetogenesis in the presence of exogenous formate in addition to H_2_ and CO_2_.

Several other *Breznakiellaceae* strains (strains 1, 4, and 6) possessed an incomplete WLP lacking the formate-activating enzymes Fhs and MetF but retaining the CODH/ACS complex (missing *acsE* in strain 1). This configuration is similar to that found in *L. azotonutricia* and is unlikely to be used for reductive acetogenesis. Instead, these genes are likely used for methionine biosynthesis, producing 5-methyl-tetrahydrofolate (THF) from acetyl-CoA and THF as described by Zhuang et al. (56). All *Breznakiellaceae* strains, including strain 2, lack formate dehydrogenase, whereas strains 2–8 encoded the formate transporter FocA. Genes associated with WLP/C1 metabolism were transcriptionally active in the fiber-associated community, supporting the physiological relevance of these C1 metabolic functions (Figure S1).

Previous metagenomic surveys reported that none of the 126 *Spirochaetota* metagenome-assembled genomes from 50 termite species encoded a complete WLP (10,57). Our results revise this view by identifying a complete, potentially functional, WLP in *Breznakiellaceae* strain 2, and incomplete WLP variants having anabolic roles in other strains. Thus, *Breznakiellaceae* likely contribute to both hydrogenotrophic acetogenesis (strain 2) and the C_1_ metabolism for amino acid biosynthesis (other strains), emphasizing metabolic diversity among strains of *Spirochaetota* in termite guts.

### Breznakiellaceae, Candidatus Fibrimonas, and Chitinispirillaceae encode multiple chitinases

Chitin is an abundant biopolymer composing the cell walls of fungi and the cuticle of insects. All reconstructed genomes encoded enzymes able to hydrolyze chitin through the combined action of chitinases (GH18, GH20, and GH23) and chitin deacetylase (CE4) (Figure 3). The genome of *Breznakiellaceae* strain 2 also contained one GH128 and one GH55, two β-1,3-glucanases that target fungal cell walls. Genes involved in chitin/NAG utilization were also transcriptionally active across multiple reconstructed genomes (Figure S1, Table S6). The presence of a complex repertoire of chitinases in the reconstructed genomes suggests that the gut microbiota of *N. takasagoensis* may digest and utilize the fungal biomass, which could serve as a nitrogen source. Moreover, these enzymes may also contribute to the recycling of the peritrophic matrix lining the midgut of termites (58–60). Gut microbes are known to provide immunity against fungal pathogens in various animals (61), and the presence of a significant number of chitinases in the genomes of *Breznakiellaceae*, *Candidatus* Fibrimonas, and *Chitinispirillaceae* raises the possibility of similar defensive functions in termites (62). Further experimental work is needed to confirm the exact role of these chitinases.

### *Breznakiellaceae, Candidatus* Fibrimonas, and *Chitinispirillaceae* fix atmospheric dinitrogen and participate in the termite nitrogen metabolism

Like the bacterial endosymbionts of many insects, the gut microbiota of termites contributes to the nitrogen metabolism of their hosts, an important function owing to the low nitrogen content of wood (1). However, the taxonomic identity of the bacteria involved in nitrogen metabolism is often obscure. We searched our 14 genomes for the genes involved in the recycling and fixation of nitrogen. Eleven strains, representing *Breznakiellaceae*, *Candidatus* Fibrimonas, and *Chitinispirillaceae*, encoded molybdenum-dependent nitrogenase genes (*nifDHK*), while the remaining three strains (two *Breznakiellaceae* and one *Candidatus* Fibrimonas) lacked a complete *nif* operon. Therefore, the ability to fix dinitrogen is strain-specific, indicating that strains of *Breznakiellaceae* and *Ca.* Fibrimonas are not functionally equivalent.

To assess the nitrogen-fixing potential of these *nif* genes, we reconstructed a phylogeny of the *nifH* sequences and found that all *nifH* genes from our genomes clustered within group I, which includes canonical nitrogenase reductases known to support nitrogen fixation. This placement suggests that the *nif* genes in our dataset are likely functional and involved in atmospheric dinitrogen reduction, unlike group IV *nifH* genes, such as those reported in *Breznakiella homolactica*, which are predicted to be involved in porphyrin metabolism rather than nitrogen fixation (31). Seven genomes of *Breznakiellaceae* also encoded vanadium-dependent nitrogenase genes (*vnfADK*), suggesting possible functional redundancy or adaptation to molybdenum-limited conditions. Expression of nitrogen fixation genes was also detected in the metatranscriptomes for genomes encoding this pathway (Figure S1, Table S6), providing independent support for the activity of the predicted nitrogen-fixation machinery.

In addition to nitrogen fixation, we examined the capacity of our 14 genomes to recycle nitrogen wastes. Only three *Breznakiellaceae* strains (strains 2, 4, and 7) encoded the complete set of urease structural genes (*ureABC*), as well as a complete urea transporter operon (*urtABCDE)* (Table S3). Other *Breznakiellaceae* strains, all *Candidatus* Fibrimonas, and *Chitinispirillaceae* lacked *ureABC*. All genomes encoded an ammonium transporter (*AmtB*), except for *Breznakiellaceae* strain 7, and all but one *Breznakiellaceae* (strain 7) and the two *Ca.* Fibrimonas strains encoded a complete glutamate biosynthesis pathway (Figures 4 and 5), suggesting a capacity to assimilate ammonia into amino acids. *Ca.* Fibrimonas 1 and *Chitinispirillaceae* 1 also possessed the oligopeptide transporter system OppABCDF, which could facilitate nitrogen acquisition from peptides, whereas OppABCDF was not identified in *Ca.* Fibrimonas 2.

Finally, the genes required for the biosynthesis of the ten amino acids essential for termites were present across our 14 genomes, supporting a nutritional symbiosis. Our results indicate that the gut *Spirochaetota* and *Fibrobacterota* of *N. takasagoensis* could play an analogous role to that of the endosymbiont *Blattabacterium* in cockroaches (63), providing nitrogenous compounds that their hosts lack from their diet. While the involvement of *Fibrobacterota* in the nitrogen metabolism of termites has previously been inferred from fragmented metagenomic contigs and lacks reliability (33), *L. azotonutricia* and *Candidatus* Treponema intracellularis, two gut *Spirochaetota* respectively associated with *Zootermopsis* and *Hodotermopsis*, together with the uncultured N□-fixer *Propulsinema mixotrichae*, an ectosymbiont of *Mixotricha paradoxa* in the gut of *Mastotermes darwiniensis*, were all known dinitrogen fixers (19,32,55,64). Our findings of functional *nif* genes in multiple strains from *N. takasagoensis* suggest that nitrogen fixation may be a general and underappreciated trait of termite-associated *Spirochaetota*. The role of *Spirochaetota* and *Fibrobacterota* in the nitrogen metabolism of *N. takasagoensis* is also reminiscent of other termite gut bacteria, such as *Candidatus* Azobacteroides, which have similar functions (9,55).

### *Breznakiellaceae, Candidatus* Fibrimonas, and *Chitinispirillaceae* are involved in other key metabolic reactions

In addition to their involvement in nitrogen metabolism, the analysis of reconstructed genomes revealed that all *Spirochaetota* and *Fibrobacterota* had other important functions. We found transporters for peptide ATP-binding proteins (*ddpDF* and *oppD*), phosphate (K14430), ferrous iron (*feoB*), cobalt (*cbiMQ* and *corA*), copper (*copA*), and thiamin (*thiQ* and *ynjD*) in multiple genomes. The numerous genes encoding alkaline phosphatase, polyphosphate kinase, and phosphate transporter in *Breznakiellaceae* genomes indicate their potential involvement in phosphate metabolism, an important function given the very low phosphorus content in wood (∼0.01%) (65).

## CONCLUSION

Termites harbor thousands of endemic bacterial species in their guts (13,14), most of which cannot be cultured (8). As an alternative to culturing, these communities are often studied using sequencing approaches; however, the metagenomes assembled with short reads are often highly fragmented, which significantly limits our ability to assign functions to specific bacteria. Here, PacBio HiFi sequencing enabled the reconstruction of highly contiguous bacterial genomes from the complex gut microbiota of *N. takasagoensis*, including multiple dominant uncultured strains. The reconstructed genomes revealed complementary roles in lignocellulose degradation, with *Fibrobacterota* primarily specializing in cellulose and *Spirochaetota* in hemicellulose degradation, together with enzymes potentially involved in lignin modification or oxidative stress tolerance. The lineages also differed in their predicted fermentation pathways and hydrogen metabolism. *Breznakiellaceae* fermented sugars to H□, CO□, acetate, and ethanol; *Ca.* Fibrimonas fermented sugars to acetate, succinate, and potentially propanoate; *Chitinispirillaceae* fermented sugars to ethanol and H. Only *Breznakiellaceae* strain 2 encoded a complete WLP potentially supporting energy-conserving reductive acetogenesis in the presence of formate, while several other strains (1, 4, and 6) possessed incomplete WLP variants likely involved in anabolic C1 metabolism. *Breznakiellaceae* and *Chitinispirillaceae* genomes also encoded the machinery for atmospheric nitrogen fixation, while the nitrogen metabolism capabilities varied among strains.

The integration of previously published metatranscriptomes from the fiber-associated community showed that key metabolic functions identified in the reconstructed genomes are transcriptionally active, providing independent support for their physiological relevance. Overall, the high contiguity of the reconstructed genomes enabled precise pathway reconstruction and revealed metabolic differences among closely related strains that are difficult to resolve from fragmented short-read MAGs. Therefore, HiFi read-based metagenomics offers an effective approach for resolving the metabolic functions of dominant uncultured members of the termite gut microbiota.

## SUPPLEMENTAL MATERIAL

The raw sequence read data together with the assembled complete bacterial genomes are available in NCBI database under project number PRJNA1126444. All additional supplementary materials are included in the supporting information.

## ACKNOWLEDGMENTS

This work was supported by the Czech Science Foundation (project No. 24-12674S) and the Japanese Society for the Promotion of Science (JSPS) (project No. 23K18035). We thank the DNA Sequencing Section and the Scientific Computation and Data Section of the Okinawa Institute of Science and Technology Graduate University, Japan, for assistance with sequencing and providing access to the OIST computing cluster Deigo, respectively.

## AUTHOR CONTRIBUTIONS

A.P. and T.B. conceptualization of the experiments. G.T. collected samples. A.P. performed the lab experiments. A.P., V.E., S.B., C.L., and Y.H. analyzed data. A.P. and T.B. wrote the paper with the significant input of G.T. and Y.H. All authors read and accepted the final version of this paper.

## MATERIALS AND METHODS

### Sample collection

We studied the gut bacterial communities of the wood-feeding termite *Nasutitermes takasagoensis*, collected on the Iriomote island (24.3301° N, 123.8188° E), Okinawa Prefecture, Japan.

### DNA extraction and sequencing

The whole guts of ten workers were dissected and submerged in PBS until the following DNA extraction with the MagAttract HMW DNA Kit, which allows the isolation of high-molecular-weight genomic DNA (100-200 kb). The library preparation was performed using the protocol for Low Input DNA (10 ng/ml) with the SMRTbell® Express Template Prep Kit 2.0 following the manufacturer’s instructions (Pacific Biosciences). The library was sequenced on the PacBio Sequel II platform using one SMRT Sequel II cell. Subreads and circular consensus sequencing (CCS) reads were generated using the SMRTLink software with default settings. A total of 251,89 Gb of raw long-read data were obtained. The Sequel II dataset was filtered after CCS correction to retain only reads of HiFi quality standards (>Q20). We obtained a total of 13.9 Gb of HiFi reads.

### Metagenome assembly and complete bacterial genome quality assessment

The quality of the HiFi reads was checked with the nanopack suite of tools (66). We assembled HiFi reads twice, once with Hifiasm-meta version 0.18.5-r499 (67) using default settings and once with Flye version 2.8.1 (68) using the options “--pacbio-hifi--meta--iterations 6”, which specifies to use six polishing iterations to assemble PacBio HiFi reads from metagenomes —with uneven coverage. The quality of the assembly was verified with QUAST v.5.0 (Quality Assessment for Genome Assemblies) (69) using the “metaquast” option.

Contigs of termite origin were identified by mapping our metagenome assemblies on a draft genome of *Nasutitermes lujae* using BLASTn 2.10.0+ (70). Contigs with hits matching termite proteins were not considered for downstream analyses. We only retained contigs matching bacterial or archaeal proteins for downstream analyses. The sequencing depth of each contig was calculated by mapping HiFi reads to the assemblies with minimap2-2.24 (71) using the parameter “-ax asm20” and SAMtools version 1.9 (72).

We used CheckM2 (73) with the option “checkm2 predict -x fasta --threads 12” to estimate the quality of each contig (completeness and contamination). This approach allowed us to identify each complete bacterial genome in our assemblies. We used simple-circularise (https://github.com/Kzra/Simple-Circularise) to circularize the linear sequences of bacterial genomes with completeness > 80%. The similarity of circular bacterial genomes obtained from Hifiasm-meta and Flye was determined using YASS (74). We only retained one copy of the genomes assembled independently by Hifiasm-meta and Flye for downstream analyses. The completeness of the retrieved genomes was estimated with BUSCO (75) using universal single-copy orthologs.

We further assessed the quality of the circular bacterial genomes by checking the consistence of sequencing depth. We mapped the reads to (1) the original bacterial genomes and (2) genome contigs joined in the 5’ and 3’ terminals and broken at the midpoint using minimap2 with options “-ax map-hifi -t 20 --secondary=no”. Sequencing depth was computed using 10 kb sliding windows with SAMtools. To estimate the representation of the reconstructed genomes in the metagenomic dataset, trimmed HiFi reads were mapped competitively against the 14 genomes using minimap2 with the “map-hifi” preset. Primary alignments with MAPQ ≥20 were retained, and the percentage of mapped reads was calculated relative to the total number of trimmed HiFi reads.

### Taxonomic assignment and functional annotation

All contigs composing the assembly were taxonomically annotated using Diamond blastx searches (76) against the Genome Taxonomy Database (GTDB) (77) with e-value of 1e-24 as a threshold. We used the lowest common ancestor (LCA) algorithm for the annotation and selected the options “--max-hsps 0 --range-culling --F 15” with output format 102. The GTDB-assigned taxonomic distribution was visualized using R and the Pavian web application (78).

Protein-coding genes were predicted using Prodigal v.2.6.3 (79). We used the option “-p meta” for the whole assembly and “-p single” for individual bacterial genomes. Protein-coding genes were functionally annotated with Prokka (80) using the command “--kingdom Bacteria”. We used Proksee (81) to assess genome features and sequence characteristics, and to visualize maps of circular genomes.

Functional annotation of protein-coding genes was performed using KAAS (KEGG Automatic Annotation Server) (82) to assign KEGG Orthology (KO) numbers, and annotation tables were also generated with DFAST (83). For genes involved in central metabolism and other key pathways, KO assignments were further checked manually by BLASTP and Conserved Domain (CD) searches. This curated annotation was used for pathway reconstruction in KEGG Mapper. Metabolic pathways were reconstructed with KofamScan (84) using E-value of 1e-10 as a threshold, which assigns KEGG Orthology (85) to protein sequences by homology searches against the Kofam database, a customized HMM database of KEGG Orthologs (KOs). KEGG annotations were used to estimate the completeness of the metabolic pathways of interest in all reconstructed genomes using KEGG-Decoder v.1.0.10 (86). Genes coding for microbial Carbohydrate-Active Enzymes (CAZymes) were identified using dbCAN (39) (dbCAN-fam-HMMs.txt.v9) and the CAZy database (40) with e-value of e-30 and minimum coverage of 0.35 set as thresholds. Hydrogenases were annotated by Hidden Markov Model (HMM) searches (87) against the PFAM database (88) accessed through the InterPro consortium (89). Catalytic sub-units of hydrogenases were classified with the HydDB tool (90).

### Phylogenetic analyses

We determined a set of bacterial marker genes for the most abundant phyla, *Spirochaetota* and *Fibrobacterota,* using CheckM2 version 1.2.1 (91), which we used to reconstruct a phylogenetic tree of all complete bacterial genomes assembled in this study and previously sequenced *Spirochaetota* and *Fibrobacterota* genomes obtained from GenBank. We aligned the proteins of each single-copy marker gene independently using MAFFT version 7.305 (92) with the parameters “--maxiterate 1000 --globalpair.” Protein alignments were concatenated with FASconCAT-G v1.11 (93). A maximum-likelihood phylogenetic tree was inferred with IQtree v. 1.6.12 (94). We used the best-fit protein mixture model “LG+F+I+G4” identified by ModelFinder (95). Branch support values were estimated with 1000 bootstraps. The tree was visualized using the R package “ggtree” (96).

### Average Nucleotide Identity (ANI) analyses

To assess genomic relatedness, pairwise Average Nucleotide Identity (ANI) comparisons were performed using FastANI v1.33 (97), a rapid alignment-free algorithm optimized for whole-genome comparisons. ANI was calculated between each of our 11 *Breznakiellaceae* and 2 *Ca.* Fibrimonas genomes, and publicly available reference genomes, including *Leadbettera azotonutricia* ZAS-9, *Treponema primitia* ZAS-2, and *Candidatus Fibrimonas termitidis* GUT221.

For the analysis, query and reference genomes in FASTA format were compared using the following parameters: fragment length of 1,000 bp and default minimum alignment fraction (20%). Genomes that did not return a result were interpreted as having ANI values below the 70% detection threshold of FastANI. Output values were used to determine species- and genus-level boundaries, with >95% ANI indicating conspecific genomes, 83–95% suggesting same-genus but different species, and <83% indicating distinct genera.

### Metatranscriptomic analyses

To evaluate whether the reconstructed genomes represent metabolically active members of the *Nasutitermes takasagoensis* gut microbiota, we analyzed previously published metatranscriptomic data from the fiber-associated bacterial community of the *N. takasagoensis* hindgut (BioProject PRJDB5939). The dataset comprises three biological replicates of paired-end Illumina RNA sequencing reads generated by Tokuda et al. (25).

The 14 reconstructed genomes were combined into a single set of genomes. Protein-coding genes were predicted and annotated using Prokka v1.14.6. Paired-end transcriptomic reads were mapped onto the genome set using Bowtie2 v2.4.5 with default parameters. Resulting SAM files were converted into BAM format, sorted, indexed using SAMtools v1.19, and filtered to retain only alignments with mapping quality ≥20.

Gene-level read counts were calculated using the program featureCounts from the Subread package v2.0.6 with paired-end mode enabled and Prokka-generated GFF annotation files. Raw counts were converted into TPM (Transcripts Per Million) values to account for differences in gene length and sequencing depth. We provided the TPM values obtained for the three biological replicates for the genes belonging to selected metabolic pathways, including cellulose degradation, hemicellulose degradation, hydrogen metabolism, Wood–Ljungdahl/C1 metabolism, nitrogen fixation and nitrogen assimilation, sulfur metabolism, chitin utilization, amino acid biosynthesis, and oxidative/redox-associated pathways. The mean pathway expression across the three biological replicates was used for visualization.

### Statistical analyses

The relative abundance of each contig was calculated by mapping the raw reads to the assembled contigs annotated as bacteria or archaea. Read counts were normalized to account for variations in sequencing depth. The relative abundance values were expressed as Transcripts per Million (TPM). The counts of individual TPM were adjusted through a centered log_2_-ratio (clr) transformation for marker genes and genes of functional significance. This adjustment was made to address the compositional structure of the data. Similarly, the contigs coverage was estimated by multiplying the mapped reads by the read length and dividing it by the contig length. Statistical analyses were performed with R version 3.4.4 (98), and data were visualized with the ggplot2 (99) package. We also visualized all genes involved in the lignocellulose digestion and the nitrogen metabolism using the R package “pheatmap” (100).

---

Figure S1. Transcriptional activity of metabolic pathways in the reconstructed *N. takasagoensis* gut bacterial genomes.

Figure S2. Sequencing depth distributions of 10 kb sliding windows for (A) the original bacterial genomes and (B) the genomes with joined terminals and breakpoint placed at the midpoint of the original contig. Dot color indicates the distance of sliding windows to 5′ terminal normalized to the length of genome.

## REFERENCES

1. Brune A. Symbiotic digestion of lignocellulose in termite guts. Nat Rev Microbiol. 2014;12(3):168–80.

2. Lynd LR, Weimer PJ, Zyl WH Van, Isak S. Microbial Cellulose Utilization: Fundamentals and Biotechnology Microbial Cellulose Utilization: Fundamentals and Biotechnology. Microbiol Mol Biol Rev. 2002;66(3):506–77.

3. Watanabe H, Noda. H. A cellulase gene of termite origin. Nature. 1998;394:330–1.

4. Watanabe H, Tokuda G. Cellulolytic systems in insects. Annu Rev Entomol. 2010;55:609–32.

5. Chouvenc T, Šobotník J, Engel MS, Bourguignon T. Termite evolution: mutualistic associations, key innovations, and the rise of Termitidae. Cell Mol Life Sci [Internet]. 2021;78(6):2749–69. Available from: 10.1007/s00018-020-03728-z

6. Hongoh Y, Sharma VK, Prakash T, Noda S, Toh H, Taylor TD, et al. Genome of an endosymbiont coupling N2 Fixation To Celluolysis Within Protist Cells in Termite Gut. 2008;(November):1108–9.

7. Potrikus CJ, Breznak JA. Gut bacteria recycle uric acid nitrogen in termites: A strategy for nutrient conservation. Proc Natl Acad Sci. 1981;78(7):4601–5.

8. Ohkuma M, Brune A. Diversity, Structure, and Evolution of the Termite Gut Microbial Community. Biol Termit A Mod Synth. 2011;413–38.

9. Hongoh Y. Toward the functional analysis of uncultivable, symbiotic microorganisms in the termite gut. Cell Mol Life Sci. 2011;68(8):1311–25.

10. Arora J, Kinjo Y, Šobotník J, Buček A, Clitheroe C, Stiblik P, et al. The functional evolution of termite gut microbiota. Microbiome [Internet]. 2022;10(1):1–22. Available from: 10.1186/s40168-022-01258-3

11. Marynowska M, Goux X, Sillam-Dussès D, Rouland-Lefèvre C, Halder R, Wilmes P, et al. Compositional and functional characterisation of biomass-degrading microbial communities in guts of plant fibre-And soil-feeding higher termites. Microbiome. 2020;8(1):1–18.

12. Warnecke F, Luginbühl P, Ivanova N, Ghassemian M, Richardson TH, Stege JT, et al. Metagenomic and functional analysis of hindgut microbiota of a wood-feeding higher termite. Nature. 2007;450(7169):560–5.

13. Bourguignon T, Lo N, Dietrich C, Šobotník J, Sidek S, Roisin Y, et al. Rampant Host Switching Shaped the Termite Gut Microbiome. Curr Biol. 2018;28(4):649–654.e2.

14. Dietrich C, Köhler T, Brune A. The cockroach origin of the termite gut microbiota: Patterns in bacterial community structure reflect major evolutionary events. Appl Environ Microbiol. 2014;80(7):2261–9.

15. Parks DH, Rinke C, Chuvochina M, Chaumeil PA, Woodcroft BJ, Evans PN, et al. Recovery of nearly 8,000 metagenome-assembled genomes substantially expands the tree of life. Nat Microbiol [Internet]. 2017;2(11):1533–42. Available from: 10.1038/s41564-017-0012-7

16. Hongoh Y, Sharma VK, Prakash T, Noda S, Taylor TD, Kudo T, et al. Complete genome of the uncultured Termite Group 1 bacteria in a single host protist cell. Proc Natl Acad Sci U S A. 2008;105(14):5555–60.

17. Kuwahara H, Yuki M, Izawa K, Ohkuma M, Hongoh Y. Genome of “Ca. Desulfovibrio trichonymphae”, an H 2-oxidizing bacterium in a tripartite symbiotic system within a protist cell in the termite gut. ISME J. 2017;11(3):766–76.

18. Takahashi K, Kuwahara H, Horikawa Y, Izawa K, Kato D, Inagaki T, et al. Emergence of putative energy parasites within Clostridia revealed by genome analysis of a novel endosymbiotic clade. ISME J. 2023;17(11):1895–906.

19. Jieyang F, Liu Y. Functional division of labor in motility, lignocellulose digestion, and nitrogen metabolism re vealed for the Mixotricha par adoxa holobiont. 2025;19(July).

20. Moss EL, Maghini DG, Bhatt AS. Complete, closed bacterial genomes from microbiomes using nanopore sequencing. Nat Biotechnol [Internet]. 2020;38(6):701–7. Available from: 10.1038/s41587-020-0422-6

21. Stewart RD, Auffret MD, Warr A, Walker AW, Roehe R, Watson M. Compendium of 4,941 rumen metagenome-assembled genomes for rumen microbiome biology and enzyme discovery. Nat Biotechnol [Internet]. 2019;37(8):953–61. Available from: 10.1038/s41587-019-0202-3

22. Kim CY, Ma J, Lee I. HiFi metagenomic sequencing enables assembly of accurate and complete genomes from human gut microbiota. Nat Commun. 2022;13(1):1–11.

23. Bickhart DM, Kolmogorov M, Tseng E, Portik DM, Korobeynikov A, Tolstoganov I, et al. Generating lineage-resolved, complete metagenome-assembled genomes from complex microbial communities. Nat Biotechnol. 2022;40(5):711–9.

24. Zhang Y, Jiang F, Yang B, Wang S, Wang H, Wang A, et al. Improved microbial genomes and gene catalog of the chicken gut from metagenomic sequencing of high-fidelity long reads. Gigascience. 2022;11:1–12.

25. Tokuda G, Mikaelyan A, Fukui C, Matsuura Y, Watanabe H, Fujishima M, et al. Fiber-associated spirochetes are major agents of hemicellulose degradation in the hindgut of wood-feeding higher termites. Proc Natl Acad Sci U S A. 2018;115(51):E11996–2004.

26. Brune A, Dietrich C. The Gut Microbiota of Termites: Digesting the Diversity in the Light of Ecology and Evolution. Annu Rev Microbiol. 2015;69(1):145–66.

27. Hongoh Y, Deevong P, Hattori S, Inoue T, Noda S, Noparatnaraporn N, et al. Phylogenetic diversity, localization, and cell morphologies of members of the candidate phylum TG3 and a subphylum in the phylum Fibrobacteres, recently discovered bacterial groups dominant in termite guts. Appl Environ Microbiol. 2006;72(10):6780–8.

28. Arora J, Buček A, Hellemans S, Beránková T, Romero Arias J, Fisher BL, et al. Evidence of cospeciation between termites and their gut bacteria on a geological time scale. Proc R Soc B Biol Sci. 2023;290(2001).

29. Brune A, Song Y, Oren A, Paster BJ. A new family for ‘termite gut treponemes’: description of Breznakiellaceae fam. nov., Gracilinema caldarium gen. nov., comb. nov., Leadbettera azotonutricia gen. nov., comb. nov., Helmutkoenigia isoptericolens gen. nov., comb. nov., and Zuelzera stenostre. Int J Syst Evol Microbiol. 2022;72(5):1–6.

30. Beránková T, Arora J, Romero Arias J, Buček A, Tokuda G, Šobotník J, et al. Termites and subsocial roaches inherited many bacterial-borne carbohydrate-active enzymes (CAZymes) from their common ancestor. Commun Biol. 2024;7(1):1449.

31. Song Y, Hervé V, Radek R, Pfeiffer F, Zheng H, Brune A. Characterization and phylogenomic analysis of Breznakiella homolactica gen. nov. sp. nov. indicate that termite gut treponemes evolved from non-acetogenic spirochetes in cockroaches. Environ Microbiol. 2021;23(8):4228–45.

32. Graber JR, Leadbetter JR, Breznak JA. Description of Treponema azotonutricium sp. nov. and Treponema primitia sp. nov., the First Spirochetes Isolated from Termite Guts. Appl Environ Microbiol. 2004;70(3):1315–20.

33. Rahman NA, Parks DH, Vanwonterghem I, Morrison M, Tyson GW, Hugenholtz P. A phylogenomic analysis of the bacterial phylum fibrobacteres. Front Microbiol. 2016;6(JAN).

34. Hungate RE. The anaerobic mesophilic cellulolytic bacteria. Bacteriol Rev. 1950;14(1):1– 49.

35. Sorokin DY, Gumerov VM, Rakitin AL, Beletsky A V., Damsté JSS, Muyzer G, et al. Genome analysis of Chitinivibrio alkaliphilus gen. nov., sp. nov., a novel extremely haloalkaliphilic anaerobic chitinolytic bacterium from the candidate phylum Termite Group 3. Environ Microbiol. 2014;16(6):1549–65.

36. Li H, Kang X, Yang M, Kasseney BD, Zhou X, Liang S, et al. Molecular insights into the evolution of woody plant decay in the gut of termites. Sci Adv. 2023;9(21).

37. Salgado JFM, Hervé V, Vera MAG, Tokuda G, Brune A. Unveiling lignocellulolytic potential: a genomic exploration of bacterial lineages within the termite gut. Microbiome [Internet]. 2024;12(1):201. Available from: 10.1186/s40168-024-01917-7

38. Graber J. BJ. Physiology and Nutrition of Treponema primitia, an H2/ CO2-Acetogenic Spirochete from Termite Hindguts. Appl Environ Microbiol. 2004;70(3):1307–14.

39. Zhang H, Yohe T, Huang L, Entwistle S, Wu P, Yang Z, et al. DbCAN2: A meta server for automated carbohydrate-active enzyme annotation. Nucleic Acids Res. 2018;46(W1):W95–101.

40. Lombard V, Ramulu HG, Drula E, Coutinho PM, Henrissat B. The carbohydrate-active enzymes database (CAZy) in 2013. 2014;42(November 2013):490–5.

41. Nargotra P, Sharma V, Lee Y-C, Tsai Y-H, Liu Y-C, Shieh C-J, et al. Microbial Lignocellulolytic Enzymes for the Effective Valorization of Lignocellulosic Biomass: A Review. Catalysts. 2022;13(1):83.

42. Suen G, Weimer PJ, Stevenson DM, Aylward FO, Boyum J, Deneke J, et al. The complete genome sequence of fibrobacter succinogenes s85 reveals a cellulolytic and metabolic specialist. PLoS One. 2011;6(4).

43. Aspeborg H, Coutinho PM, Wang Y, Brumer H, Henrissat B. Evolution, substrate specificity and subfamily classification of glycoside hydrolase family 5 (GH5). BMC Evol Biol [Internet]. 2012;12(1):1. Available from: BMC Evolutionary Biology

44. Allardyce BJ, Linton SM, Saborowski R. The last piece in the cellulase puzzle: The characterisation of β-glucosidase from the herbivorous gecarcinid land crab Gecarcoidea natalis. J Exp Biol. 2010;213(17):2950–7.

45. Weimer PJ. Effects of dilution rate and pH on the ruminal cellulolytic bacterium Fibrobacter succinogenes S85 in cellulose-fed continuous culture. Arch Microbiol. 1993;160(4):288–94.

46. Glasgow EM, Kemna EI, Bingman CA, Ing N, Deng K, Bianchetti CM, et al. A structural and kinetic survey of GH5_4 endoglucanases reveals determinants of broad substrate specificity and opportunities for biomass hydrolysis. J Biol Chem. 2020;295(51):17752– 69.

47. Albersheim, P., Darvill, A., Roberts, K., Sederoff, R., & Staehelin A. Plant Cell Walls. Garl Sci [Internet]. 2010; Available from: 10.1201/9780203833476

48. Odelson DA, Breznak JA. Volatile fatty acid production by the hindgut microbiota of xylophagous termites. Appl Environ Microbiol. 1983;45(5):1602–13.

49. Kwong WK, Zheng H, Moran NA. Convergent evolution of a modified, acetate-driven TCA cycle in bacteria. Nat Microbiol. 2017;2(April):10–2.

50. Hackmann TJ, Ngugi DK, Firkins JL, Tao J. Genomes of rumen bacteria encode atypical pathways for fermenting hexoses to short-chain fatty acids. Environ Microbiol. 2017;19(11):4670–83.

51. Ballor NR, Paulsen I, Leadbetter JR. Genomic Analysis Reveals Multiple [FeFe] Hydrogenases and Hydrogen Sensors Encoded by Treponemes from the H 2-Rich Termite Gut. Microb Ecol. 2012;63(2):282–94.

52. Brauman A, Kane MD, Labat M, Breznak JA. Genesis of Acetate and Methane by Gut Bacteria of Nutritionally Diverse Termites. Science (80-). 1992;257(5075):1384–7.

53. Pester M, Brune A. Hydrogen is the central free intermediate during lignocellulose degradation by termite gut symbionts. ISME J. 2007;1(6):551–65.

54. Breznak JA, Switzer JM. Acetate synthesis from H2 plus CO2 by termite gut microbes. Appl Environ Microbiol. 1986;52(4):623–30.

55. Ohkuma M, Noda S, Hattori S, Iida T, Yuki M, Starns D, et al. Acetogenesis from H2 plus CO2and nitrogen fixation by an endosymbiotic spirochete of a termite-gut cellulolytic protist. Proc Natl Acad Sci U S A. 2015;112(33):10224–30.

56. Zhuang WQ, Yi S, Bill M, Brisson VL, Feng X, Men Y, et al. Incomplete Wood-Ljungdahl pathway facilitates one-carbon metabolism in organohalide-respiring Dehalococcoides mccartyi. Proc Natl Acad Sci U S A. 2014;111(17):6419–24.

57. Hervé V, Liu P, Dietrich C, Sillam-Dussès D, Stiblik P, Šobotník J, et al. Phylogenomic analysis of 589 metagenome-assembled genomes encompassing all major prokaryotic lineages from the gut of higher termites. PeerJ. 2020;2020(2):1–27.

58. Fujita A, Hojo M, Aoyagi T, Hayashi Y, Arakawa G, Tokuda G, et al. Details of the digestive system in the midgut of Coptotermes formosanus Shiraki. J Wood Sci. 2010;56(3):222–6.

59. A. Morales-Ramos M. G. R and H. S-B. Peritrophic Matrix of the Formosan Subterranean Termite (Isoptera: Rhinotermitidae). Florida Entomol Soc [Internet]. 2006;89(1):99–101. Available from: https://www.jstor.org/stable/3496

60. Sandoval Mojica, A. F., & Scharf ME. Gut genes associated with the peritrophic matrix in Reticulitermes flavipes (Blattodea: Rhinotermitidae): Identification and characterization. Arch Insect Biochem Physiol. 2016;(92(2)):127–42.

61. Boucias DG, Zhou Y, Huang S, Keyhani NO. Microbiota in insect fungal pathology. Appl Microbiol Biotechnol. 2018;102(14):5873–88.

62. Nishimura Y, Otagiri M, Yuki M, Shimizu M, Inoue J ichi, Moriya S, et al. Division of functional roles for termite gut protists revealed by single-cell transcriptomes. ISME J [Internet]. 2020;14(10):2449–60. Available from: 10.1038/s41396-020-0698-z

63. Sabree ZL, Kambhampati S, Moran NA. Nitrogen recycling and nutritional provisioning by Blattabacterium, the cockroach endosymbiont. Proc Natl Acad Sci U S A. 2009;106(46):19521–6.

64. Lilburn TG, Kim KS, Ostrom NE, Byzek KR, Leadbetter JR, Breznak JA. Nitrogen fixation by symbiotic and free-living spirochetes. Science (80-). 2001;292(5526):2495–8.

65. Antikainen R, Haapanen R, Rekolainen S. Flows of nitrogen and phosphorus in Finland - The forest industry and use of wood fuels. J Clean Prod. 2004;12(8–10):919–34.

66. De Coster W, D’Hert S, Schultz DT, Cruts M, Van Broeckhoven C. NanoPack: Visualizing and processing long-read sequencing data. Bioinformatics. 2018;34(15):2666– 9.

67. Cheng H, Concepcion GT, Feng X, Zhang H, Li H. Haplotype-resolved de novo assembly using phased assembly graphs with hifiasm. Nat Methods [Internet]. 2021;18(2):170–5. Available from: 10.1038/s41592-020-01056-5

68. Kolmogorov M, Yuan J, Lin Y, Pevzner PA. Assembly of long, error-prone reads using repeat graphs. Nat Biotechnol [Internet]. 2019;37(5):540–6. Available from: 10.1038/s41587-019-0072-8

69. Gurevich A, Saveliev V, Vyahhi N, Tesler G. QUAST: Quality assessment tool for genome assemblies. Bioinformatics. 2013;29(8):1072–5.

70. Altschul SF, Gish W, Miller W, Myers EW, Lipman DJ. Basic local alignment search tool. J Mol Biol. 1990;215(3):403–10.

71. Heng L. Minimap2: Pairwise alignment for nucleotide sequences. Bioinformatics. 2018;34(18):3094–100.

72. Danecek P, Bonfield JK, Liddle J, Marshall J, Ohan V, Pollard MO, et al. Twelve years of SAMtools and BCFtools. Gigascience. 2021;10(2):1–4.

73. Chklovski, A., Parks, D.H., Woodcroft BJ et al. CheckM2: a rapid, scalable and accurate tool for assessing microbial genome quality using machine learning. Nat Methods. 2023;20:1203–12.

74. Noé L, Kucherov G. YASS: Enhancing the sensitivity of DNA similarity search. Nucleic Acids Res. 2005;33(SUPPL. 2):540–3.

75. Simão FA, Waterhouse RM, Ioannidis P, Kriventseva E V., Zdobnov EM. BUSCO: Assessing genome assembly and annotation completeness with single-copy orthologs. Bioinformatics. 2015;31(19):3210–2.

76. Buchfink B, Reuter K, Drost HG. Sensitive protein alignments at tree-of-life scale using DIAMOND. Nat Methods [Internet]. 2021;18(4):366–8. Available from: 10.1038/s41592-021-01101-x

77. Parks DH, Chuvochina M, Rinke C, Mussig AJ, Chaumeil PA, Hugenholtz P. GTDB: An ongoing census of bacterial and archaeal diversity through a phylogenetically consistent, rank normalized and complete genome-based taxonomy. Nucleic Acids Res. 2022;50(D1):D785–94.

78. Breitwieser FP, Salzberg SL. Pavian: Interactive analysis of metagenomics data for microbiome studies and pathogen identification. Bioinformatics. 2020;36(4):1303–4.

79. Hyatt D, Chen G-L, LoCascio PF, Land ML, Larimer FW, Hauser LJ. Prodigal: prokaryotic gene recognition and translation initiation site identification. Nat Commun [Internet]. 2010;6(1):1–8. Available from: http://dx.doi.org/10.1016/B978-0-12-407863-5.00023-X%5Cnhttp://www.nature.com/doifinder/10.1038/ismej.2009.79%5Cnhttp://www.nature. com/doifinder/10.1038/nature09916%5Cnhttp://dx.doi.org/10.1038/srep25982%5Cnhttp//:dx.doi.org/10.1038/ismej.2010.144%5Cnhttp

80. Seemann T. Prokka: Rapid prokaryotic genome annotation. Bioinformatics. 2014;30(14):2068–9.

81. Grant JR, Enns E, Marinier E, Mandal A, Herman EK, Chen CY, et al. Proksee: In-depth characterization and visualization of bacterial genomes. Nucleic Acids Res. 2023;51(W1):W484–92.

82. Moriya Y, Itoh M, Okuda S, Yoshizawa AC, Kanehisa M. KAAS: An automatic genome annotation and pathway reconstruction server. Nucleic Acids Res. 2007;35(SUPPL.2):182–5.

83. Tanizawa Y, Fujisawa T, Nakamura Y. DFAST: A flexible prokaryotic genome annotation pipeline for faster genome publication. Bioinformatics. 2018;34(6):1037–9.

84. Aramaki T, Blanc-Mathieu R, Endo H, Ohkubo K, Kanehisa M, Goto S, et al. KofamKOALA: KEGG Ortholog assignment based on profile HMM and adaptive score threshold. Bioinformatics. 2020;36(7):2251–2.

85. Kanehisa M, Goto S. KEGG: Kyoto Encyclopedia of Genes and Genomes. Nucleic Acids Res. 2000;28(1):27–30.

86. Graham ED, Heidelberg JF, Tully BJ. Potential for primary productivity in a globally-distributed bacterial phototroph. ISME J [Internet]. 2018;12(7):1861–6. Available from: 10.1038/s41396-018-0091-3

87. Johnson LS, Eddy SR, Portugaly E. Johnson, L.S.; Eddy, S.R.; Portugaly, E. Hidden Markov model speed heuristic and iterative HMM search procedure. BMC Bioinform. 2010, 11, 431. BMC Bioinformatics. 2010;11:431.

88. Mistry J, Chuguransky S, Williams L, Qureshi M, Salazar GA, Sonnhammer ELL, et al. Pfam: The protein families database in 2021. Nucleic Acids Res. 2021;49(D1):D412–9.

89. Blum M, Chang HY, Chuguransky S, Grego T, Kandasaamy S, Mitchell A, et al. The InterPro protein families and domains database: 20 years on. Nucleic Acids Res. 2021;49(D1):D344–54.

90. Søndergaard D, Pedersen CNS, Greening C. HydDB: A web tool for hydrogenase classification and analysis. Nat Publ Gr. 2016;1–8.

91. Parks DH, Imelfort M, Skennerton CT, Hugenholtz P, Tyson GW. CheckM: Assessing the quality of microbial genomes recovered from isolates, single cells, and metagenomes. Genome Res. 2015;25(7):1043–55.

92. Katoh K, Standley DM. MAFFT multiple sequence alignment software version 7: Improvements in performance and usability. Mol Biol Evol. 2013;30(4):772–80.

93. Kück P, Longo GC. FASconCAT-G: Extensive functions for multiple sequence alignment preparations concerning phylogenetic studies. Front Zool. 2014;11(1):1–8.

94. Nguyen LT, Schmidt HA, Von Haeseler A, Minh BQ. IQ-TREE: A fast and effective stochastic algorithm for estimating maximum-likelihood phylogenies. Mol Biol Evol. 2015;32(1):268–74.

95. Kalyaanamoorthy S, Minh BQ, Wong TKF, Von Haeseler A, Jermiin LS. ModelFinder: Fast model selection for accurate phylogenetic estimates. Nat Methods. 2017;14(6):587–9.

96. Yu G, Smith DK, Zhu H, Guan Y, Lam TTY. Ggtree: an R Package for Visualization and Annotation of Phylogenetic Trees With Their Covariates and Other Associated Data. Methods Ecol Evol. 2017;8(1):28–36.

97. Jain C, Rodriguez-R LM, Phillippy AM, Konstantinidis KT, Aluru S. High throughput ANI analysis of 90K prokaryotic genomes reveals clear species boundaries. Nat Commun. 2018;9(1):1–8.

98. R Core Team. R: A language and environment for statistical computing. R Found Stat Comput Vienna, Austria. 2021;

99. Wickham H. ggplot2: Elegant Graphics for Data Analysis. Springer-Verlag New York [Internet]. Vol. 35, Media. 2016. 211 p. Available from: http://link.springer.com/10.1007/978-0-387-98141-3

100. Kolde R. Package “pheatmap”: Pretty heatmaps. R Packag. 2022;1–8.

