## Supplementary figures and images for "High-fidelity long-read metagenomics reveals the metabolic functions of dominant termite gut bacteria"

### Supplemental figure S1

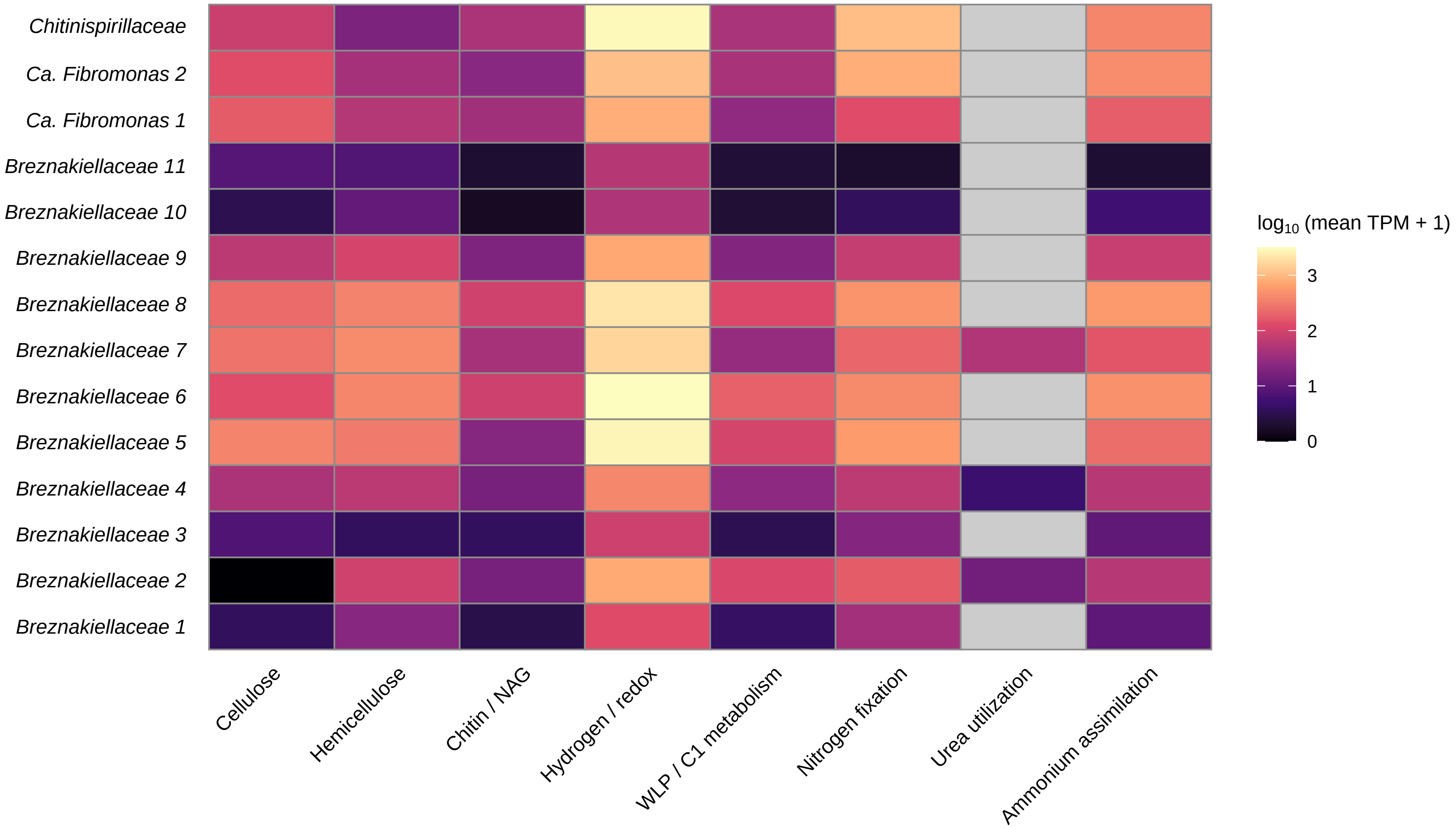

### Supplemental figure S2

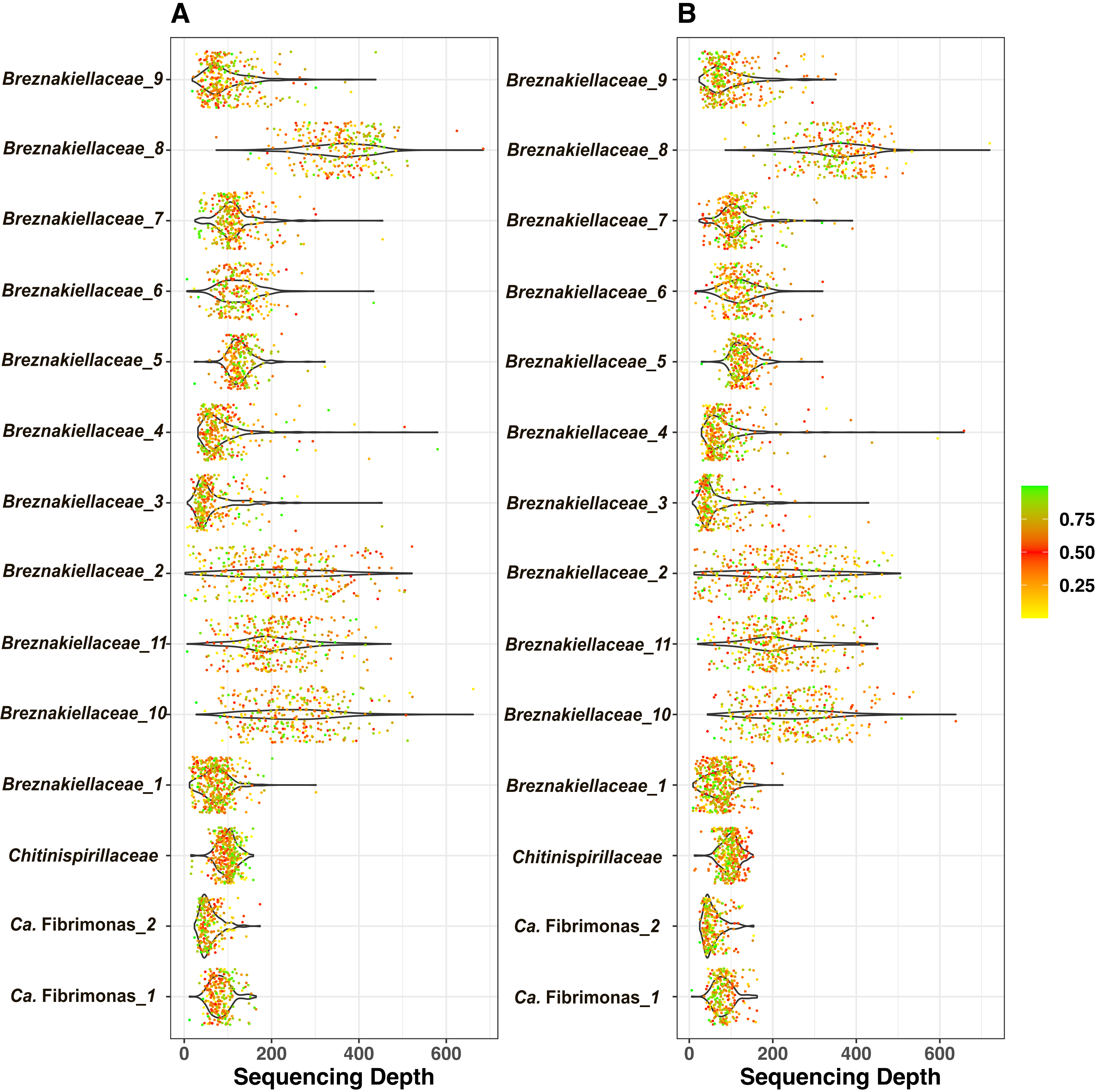
