## Supplemental table S1 for "High-fidelity long-read metagenomics reveals the metabolic functions of dominant termite gut bacteria"

| GENOMES | Completeness [%] | Contamination [%] | GC [%] | Coding density [%] | rRNA | tRNA | CDCs | Genome size [Mbp] |
|---|---|---|---|---|---|---|---|---|
| *Breznakiellaceae* strain 1 | 100 | 0.71 | 46 | 88.59 | 6 | 47 | 4217 | 4.24 |
| *Breznakiellaceae* strain 2 | 98.4 | 2.44 | 48 | 90.5 | 6 | 45 | 3109 | 3.33 |
| *Breznakiellaceae* strain 3 | 94.15 | 1.4 | 48 | 91.92 | 6 | 46 | 2540 | 2.62 |
| *Breznakiellaceae* strain 4 | 99.3 | 0.9 | 40 | 92.58 | 3 | 49 | 3160 | 2.65 |
| *Breznakiellaceae* strain 5 | 100 | 0.86 | 40 | 90.85 | 3 | 46 | 2522 | 3.25 |
| Breznakiellaceae strain 6 | 99.26 | 4.8 | 41 | 90.21 | 3 | 48 | 2478 | 3.23 |
| *Breznakiellaceae* strain 7 | 96.55 | 3.69 | 43 | 92.3 | 6 | 47 | 3110 | 2.79 |
| *Breznakiellaceae* strain 8 | 98.31 | 0.09 | 45 | 90.64 | 6 | 47 | 2593 | 3.27 |
| *Breznakiellaceae* strain 9 | 98.3 | 1.91 | 45 | 90.82 | 6 | 46 | 3035 | 3.2 |
| *Breznakiellaceae* strain 10 | 97.33 | 2.07 | 44 | 90.78 | 6 | 47 | 2949 | 3.19 |
| *Breznakiellaceae* strain 11 | 100 | 0.62 | 45 | 91.01 | 6 | 46 | 2897 | 2.67 |
| *Ca.* Fibrimonas strain 1 | 97.6 | 0.58 | 44 | 94.5 | 4 | 45 | 2558 | 2.52 |
| *Ca.* Fibrimonas strain 2 | 79.85 | 6.86 | 43 | 94.12 | 2 | 35 | 2403 | 2.3 |
| *Chitinispirillaceae* strain 1 | 99.01 | 0.4 | 54 | 86.93 | 6 | 46 | 3232 | 3.65 |
