## Supplemental table S3 for "High-fidelity long-read metagenomics reveals the metabolic functions of dominant termite gut bacteria"

Supplementary table S3. Annotation details of the genes encoding metabolic pathways of the reconstructed genomes. The genes are sorted by functional groups (in blue) and subgroups (in grey). "0" shows that gene is absent, "1" shows that gene is present.

| Gene name | Gene product | KO term | Enzymes | *Breznakiellaceae* strain 1 | *Breznakiellaceae* strain 2 | *Breznakiellaceae* strain 3 | *Breznakiellaceae* strain 4 | *Breznakiellaceae* strain 5 | *Breznakiellaceae* strain 6 | *Breznakiellaceae* strain 7 | *Breznakiellaceae* strain 8 | *Breznakiellaceae* strain 9 | *Breznakiellaceae* strain 10 | *Breznakiellaceae* strain 11 | Ca. Fibrimonas strain 1 | Ca. Fibrimonas strain 2 | Chitinispirillaceae strain 1 |
|---|---|---|---|---|---|---|---|---|---|---|---|---|---|---|---|---|---|
| **Glycolysis** | | | | | | | | | | | | | | | | | |
| **Embden-Meyerhof pathway** | | | | | | | | | | | | | | | | | |
| *pgm* | Phosphoglucomutase | K01835 | EC:5.4.2.2 | 1 | 1 | 1 | 1 | 1 | 1 | 1 | 1 | 1 | 1 | 1 | 1 | 1 | 1 |
| *pgi* | Glucose-6-phosphate isomerase | K01810 | EC:5.3.1.9 | 1 | 1 | 1 | 1 | 1 | 1 | 1 | 1 | 1 | 1 | 1 | 1 | 1 | 1 |
| *ALDO* | Fructose-bisphosphate aldolase class I | K01623 | EC:4.1.2.13 | 1 | 1 | 1 | 1 | 1 | 1 | 1 | 1 | 1 | 1 | 1 | 1 | 1 | 1 |
| *pgk* | 3-Phosphoglycerate kinase | K00927 | EC:2.7.2.3 | 1 | 1 | 1 | 1 | 1 | 1 | 1 | 1 | 1 | 1 | 1 | 1 | 1 | 1 |
| *gpmI* | 2,3-Bisphosphoglycerate-independent phosphoglycerate mutase | K15633 | EC:5.4.2.12 | 1 | 1 | 0 | 1 | 1 | 1 | 1 | 1 | 1 | 1 | 1 | 1 | 1 | 1 |
| *eno* | Enolase/phosphopyruvate hydratase | K01689 | EC:4.2.1.11 | 1 | 1 | 0 | 1 | 1 | 1 | 1 | 1 | 1 | 1 | 1 | 1 | 1 | 0 |
| *pyk* | Pyruvate kinase | K00873 | EC:2.7.1.40 | 1 | 1 | 1 | 1 | 1 | 1 | 1 | 1 | 1 | 1 | 1 | 1 | 1 | 1 |
| **Entner-Doudoroff pathway** | | | | | | | | | | | | | | | | | |
| *zwf* | glucose-6-phosphate 1-dehydrogenase | K00036 | EC:1.1.1.49 | 1 | 0 | 0 | 0 | 0 | 0 | 0 | 0 | 0 | 0 | 0 | 0 | 0 | 0 |
| *pgl* | 6-Phosphogluconolactonase | K07404 | EC:3.1.1.31 | 1 | 1 | 1 | 1 | 1 | 1 | 1 | 1 | 1 | 1 | 1 | 0 | 0 | 0 |
| *edd* | Phosphogluconate dehydratase | K01690 | EC:4.2.1.12 | 1 | 1 | 1 | 1 | 1 | 1 | 1 | 1 | 1 | 1 | 1 | 1 | 0 | 1 |
| *eda* | 2-dehydro-3-deoxyphosphogluconate aldolase | K01625 | EC:4.1.2.14 | 1 | 1 | 1 | 1 | 1 | 1 | 1 | 1 | 1 | 1 | 1 | 1 | 1 | 1 |
| *gnaD* | Gluconate/galactonate dehydratase | K05308 | EC:4.2.1.140 | 1 | 1 | 0 | 1 | 1 | 1 | 1 | 1 | 1 | 1 | 1 | 0 | 0 | 0 |
| *kdgK* | 2-dehydro-3-deoxygluconokinase | K00874 | EC:2.7.1.45 | 1 | 1 | 1 | 1 | 1 | 1 | 1 | 1 | 1 | 1 | 1 | 1 | 1 | 1 |
| *gapN* | Glyceraldehyde-3-phosphate dehydrogenase (NADP+) | K00131 | EC:1.2.1.9 | 1 | 1 | 1 | 1 | 1 | 1 | 1 | 1 | 1 | 1 | 1 | 1 | 1 | 1 |
| **Pentose phosphate pathway** | | | | | | | | | | | | | | | | | |
| *tktA, tktB* | Transketolase | K00615 | EC:2.2.1.1 | 1 | 0 | 1 | 0 | 0 | 0 | 0 | 0 | 1 | 0 | 0 | 0 | 0 | 0 |
| *talA, talB* | Transaldolase | K00616 | EC:2.2.1.2 | 1 | 0 | 1 | 1 | 1 | 1 | 1 | 1 | 1 | 1 | 1 | 0 | 0 | 0 |
| *rpe* | Ribulose-5-phosphate 3-epimerase | K01783 | EC:5.1.3.1 | 1 | 1 | 1 | 1 | 1 | 1 | 1 | 1 | 1 | 1 | 1 | 1 | 1 | 1 |
| *rpiAB* | Ribose-5-phosphate isomerase | K01807 | EC:5.3.1.6 | 1 | 1 | 1 | 1 | 1 | 1 | 1 | 1 | 1 | 1 | 1 | 1 | 1 | 1 |
| **Wood–Ljungdahl pathway** | | | | | | | | | | | | | | | | | |
| *fdhA* | Formate dehydrogenase (NADP+) alpha subunit | K05299 | EC:1.17.1.10 | 1 | 1 | 1 | 1 | 1 | 1 | 1 | 1 | 1 | 1 | 1 | 1 | 1 | 1 |
| *fdhB* | Formate dehydrogenase (NADP+) beta subunit | K15022 | EC:1.17.1.10 | 1 | 1 | 1 | 1 | 1 | 1 | 1 | 1 | 1 | 1 | 1 | 1 | 1 | 1 |
| *fdhF* | Formate dehydrogenase (hydrogenase) | K22015 | EC:1.17.98.4 | 0 | 0 | 0 | 0 | 0 | 0 | 0 | 0 | 0 | 0 | 0 | 0 | 0 | 0 |
| *fhs* | Formyl-THF synthetase/Formate-THF ligase | K01938 | EC:6.3.4.3 | 0 | 1 | 1 | 1 | 1 | 1 | 1 | 1 | 1 | 1 | 1 | 0 | 0 | 1 |
| *folD* | Formyl-THF cyclohydrolase&Methylene-THF dehydrogenase | K01491 | EC:3.5.4.9 | 1 | 1 | 1 | 1 | 1 | 1 | 1 | 1 | 1 | 1 | 1 | 1 | 1 | 1 |
| *metF* | Methylene-THF reductase | K00297 | EC:1.5.1.20 | 1 | 1 | 1 | 0 | 1 | 0 | 1 | 1 | 1 | 1 | 1 | 1 | 1 | 1 |
| *acsACDE* | 5-methyltetrahydrofolate corrinoid | K15023, | EC:2.1.1.258 | 1 | 1 | 1 | 1 | 1 | 1 | 1 | 1 | 1 | 1 | 1 | 1 | 1 | 0 |
| **Phosphate acetyltransferase-acetate kinase pathway** | | | | | | | | | | | | | | | | | |
| *pta* | Phosphate acetyltransferase | K00625 | EC:2.3.1.8 | 1 | 1 | 1 | 1 | 1 | 1 | 1 | 1 | 1 | 1 | 1 | 1 | 1 | 1 |
| *ackA* | Acetate kinase | K00925 | EC:2.7.2.1 | 1 | 1 | 1 | 1 | 1 | 1 | 1 | 1 | 1 | 1 | 1 | 1 | 0 | 0 |
| **Chitin degradation** | | | | | | | | | | | | | | | | | |
| | Chitinase | K01183 | EC:3.2.1.14 | 1 | 1 | 1 | 1 | 1 | 1 | 1 | 1 | 1 | 1 | 1 | 1 | 1 | 0 |
| *nagZ* | Beta-N-acetylhexosaminidase | K01207 | EC:3.2.1.52 | 1 | 1 | 1 | 1 | 1 | 1 | 1 | 1 | 1 | 1 | 1 | 0 | 0 | 0 |
| *NAGK* | N-acetylglucosamine kinase | K00884 | EC:2.7.1.59 | 1 | 1 | 0 | 1 | 1 | 1 | 1 | 1 | 0 | 1 | 1 | 1 | 1 | 1 |
| *nagA* | N-acetylglucosamine-6-phosphate deacetylase | K01443 | EC:3.5.1.25 | 1 | 1 | 0 | 1 | 1 | 1 | 1 | 1 | 1 | 1 | 1 | 0 | 0 | 1 |
| *nagB* | Glucosamine-6-phosphate deaminase | K02564 | EC:3.5.99.6 | 1 | 1 | 0 | 1 | 1 | 0 | 0 | 0 | 0 | 0 | 0 | 0 | 0 | 0 |
| **Glycogen metabolism** | | | | | | | | | | | | | | | | | |
| *glgC* | Glucose-1-phosphate adenylyltransferase | K00975 | EC:2.7.7.27 | 1 | 1 | 1 | 1 | 1 | 1 | 1 | 1 | 1 | 1 | 1 | 1 | 0 | 1 |
| *glgA* | Glycogen synthase | K00703 | EC:2.4.1.21 | 1 | 1 | 1 | 1 | 1 | 1 | 1 | 1 | 1 | 1 | 1 | 1 | 1 | 1 |
| *glgB* | 1,4-Alpha-glucan branching enzyme | K00700 | EC:2.4.1.18 | 1 | 1 | 1 | 1 | 1 | 1 | 1 | 1 | 1 | 1 | 1 | 1 | 1 | 1 |
| *glgP* | Glucan phosphorylase | K00688 | EC:2.4.1.1 | 1 | 1 | 1 | 1 | 1 | 1 | 1 | 1 | 1 | 1 | 1 | 1 | 1 | 1 |
| *treX, glgX* | Pullulanase/glycogen debranching enzyme | K02438 | EC:3.2.1. | 1 | 1 | 1 | 1 | 1 | 1 | 1 | 1 | 1 | 1 | 1 | 0 | 0 | 0 |
| **Pyruvate metabolism** | | | | | | | | | | | | | | | | | |
| *porC* | Pyruvate ferredoxin oxidoreductase gamma subunit | K00172 | EC:1.2.7.1 | 1 | 1 | 1 | 1 | 1 | 1 | 1 | 1 | 1 | 1 | 1 | 1 | 1 | 1 |

Supplementary table S3. Annotation details of the genes encoding metabolic pathways of the reconstructed genomes. The genes are sorted by functional groups (in blue) and subgroups (in grey). "0" shows that gene is absent, "1" shows that gene is present.

| | | | | | | | | | | | | | | | | | |
|---|---|---|---|---|---|---|---|---|---|---|---|---|---|---|---|---|---|
| **TCA cycle** | | | | | | | | | | | | | | | | | |
| *sfcA, maeA* | Malic enzyme/malate dehydrogenase (oxaloacetate-decarboxyla | K00027 | EC:1.1.1.38 | 1 | 1 | 1 | 1 | 1 | 1 | 1 | 1 | 1 | 1 | 1 | 1 | 0 | 1 |
| *gltA* | Citrate synthase | K01647 | EC:2.3.3.1 | 0 | 1 | 1 | 1 | 1 | 1 | 1 | 1 | 1 | 1 | 1 | 1 | 0 | 1 |
| *acnA* | Aconitase/aconitate hydratase | K01681 | EC:4.2.1.3 | 1 | 1 | 1 | 1 | 1 | 1 | 1 | 1 | 1 | 1 | 1 | 1 | 1 | 1 |
| *icd* | Isocitrate dehydrogenase | K00031 | EC:1.1.1.42 | 1 | 1 | 1 | 1 | 1 | 1 | 1 | 1 | 1 | 1 | 1 | 1 | 1 | 1 |
| **2-Oxoglutarate ferredoxin oxidoreductase complex (OFOR)** | | | | | | | | | | | | | | | | | |
| *korD, oorD* | 2-Oxoglutarate ferredoxin oxidoreductase subunit delta | K00176 | EC:1.2.7.3 | 1 | 1 | 1 | 1 | 1 | 1 | 1 | 1 | 1 | 1 | 1 | 1 | 1 | 1 |
| *korA, oorA, (* | 2-Oxoglutarate ferredoxin oxidoreductase subunit alpha | K00174 | EC:1.2.7.3 | 1 | 1 | 1 | 1 | 1 | 1 | 1 | 1 | 1 | 1 | 1 | 1 | 1 | 1 |
| *korB, oorB, (* | 2-Oxoglutarate ferredoxin oxidoreductase subunit beta | K00175 | EC:1.2.7.11 | 1 | 1 | 1 | 1 | 1 | 1 | 1 | 1 | 1 | 1 | 1 | 1 | 1 | 1 |
| *korC, oorC* | 2-Oxoglutarate ferredoxin oxidoreductase subunit gamma | K00177 | EC:1.2.7.3 | 1 | 1 | 1 | 1 | 1 | 1 | 1 | 1 | 1 | 1 | 1 | 1 | 0 | 1 |
| *fumA, fumB* | Fumarate hydratase, class I | K01676 | EC:4.2.1.2 | 1 | 1 | 1 | 1 | 1 | 1 | 1 | 1 | 1 | 1 | 1 | 1 | 1 | 1 |
| **Succinate dehydrogenase/fumarate reductase** | | | | | | | | | | | | | | | | | |
| *frdA* | Succinate dehydrogenase flavoprotein subunit | K00244 | EC:1.3.5.4 | 1 | 1 | 1 | 1 | 1 | 1 | 1 | 1 | 1 | 1 | 1 | 1 | 1 | 1 |
| *sdhB, frdB* | Succinate dehydrogenase/fumarate reductase, Fe-S protein subu | K00240 | EC:1.3.5.1 | 1 | 1 | 1 | 1 | 1 | 1 | 1 | 1 | 1 | 1 | 1 | 1 | 1 | 1 |
| *urdA* | Urocanate reductase | K17363 | EC:1.3.99.33 | 1 | 1 | 1 | 1 | 1 | 1 | 1 | 1 | 1 | 1 | 1 | 1 | 1 | 1 |
| *nadB* | L-aspartate oxidase | K00278 | EC:1.4.3.16 | 1 | 1 | 1 | 1 | 1 | 1 | 1 | 1 | 1 | 1 | 1 | 1 | 1 | 1 |
| **Fermentation** | | | | | | | | | | | | | | | | | |
| **Lactate** | | | | | | | | | | | | | | | | | |
| *ldhA* | D-lactate dehydrogenase | K03778 | EC:1.1.1.28 | 1 | 1 | 1 | 1 | 1 | 1 | 1 | 1 | 1 | 1 | 1 | 1 | 1 | 1 |
| *ldh* | L-lactate dehydrogenase | K00016 | EC:1.1.1.27 | 0 | 0 | 0 | 0 | 0 | 0 | 0 | 0 | 0 | 0 | 0 | 1 | 1 | 0 |
| | Lactate 2-monooxygenase | K00467 | EC:1.13.12.4 | 1 | 1 | 1 | 1 | 1 | 1 | 1 | 1 | 1 | 1 | 1 | 1 | 1 | 1 |
| *larA* | Lactate racemase | K22373 | EC:5.1.2.1 | 1 | 1 | 1 | 1 | 1 | 1 | 1 | 1 | 1 | 1 | 1 | 0 | 0 | 1 |
| **Acetate** | | | | | | | | | | | | | | | | | |
| *pta* | Phosphate acetyltransferase | K00625 | EC:2.3.1.8 | 1 | 1 | 1 | 1 | 1 | 1 | 1 | 1 | 1 | 1 | 1 | 1 | 1 | 1 |
| *ackA* | Acetate kinase | K00925 | EC:2.7.2.1 | 1 | 1 | 1 | 1 | 1 | 1 | 1 | 1 | 1 | 1 | 1 | 1 | 0 | 0 |
| **Ethanol** | | | | | | | | | | | | | | | | | |
| *adhE* | Acetaldehyde dehydrogenase / alcohol dehydrogenase | K04072 | EC:1.2.1.10 | 1 | 1 | 1 | 1 | 1 | 1 | 1 | 1 | 1 | 1 | 1 | 1 | 1 | 1 |
| *ALDH* | Aldehyde dehydrogenase (NAD+) | K00128 | EC:1.2.1.3 | 1 | 1 | 1 | 1 | 1 | 1 | 1 | 1 | 1 | 1 | 1 | 1 | 1 | 1 |
| **Alanine** | | | | | | | | | | | | | | | | | |
| *AGXT2* | Alanine-glyoxylate transaminase / (R)-3-amino-2-methylpropion | K00827 | EC:2.6.1.44 | 1 | 1 | 1 | 1 | 1 | 1 | 1 | 1 | 1 | 1 | 1 | 1 | 1 | 1 |
| *ald* | Alanine dehydrogenase | K00259 | EC:1.4.1.1 | 1 | 1 | 1 | 1 | 1 | 1 | 1 | 1 | 1 | 1 | 1 | 1 | 1 | 1 |
| **Cellobiose** | | | | | | | | | | | | | | | | | |
| | Cellobiose phosphorylase | K00702 | EC:2.4.1.20 | 1 | 1 | 1 | 1 | 1 | 1 | 1 | 1 | 1 | 1 | 1 | 1 | 0 | 1 |
| *glk* | Glucokinase | K00845 | EC:2.7.1.2 | 1 | 1 | 1 | 1 | 1 | 1 | 1 | 1 | 1 | 1 | 1 | 1 | 1 | 1 |
| **Hydrogenase** | | | | | | | | | | | | | | | | | |
| *hydA* | Sulfhydrogenase subunit alpha | K17993 | EC:1.12.1.3 | 0 | 0 | 0 | 0 | 0 | 0 | 0 | 0 | 0 | 0 | 0 | 1 | 1 | 0 |
| *hydA2* | Iron hydrogenase HydA2 | K25123 | EC:1.12.7.- | 1 | 1 | 1 | 1 | 1 | 1 | 1 | 1 | 1 | 1 | 1 | 1 | 1 | 1 |
| *hndA* | NADP-reducing hydrogenase subunit HndA | K18330 | EC:1.12.1.3 | 1 | 1 | 1 | 1 | 1 | 1 | 1 | 1 | 1 | 1 | 1 | 1 | 1 | 1 |
| *hndB* | NADP-reducing hydrogenase subunit HndB | K17992 | EC:1.12.1.3 | 1 | 1 | 1 | 1 | 1 | 1 | 1 | 1 | 1 | 1 | 1 | 1 | 1 | 1 |
| *hndC* | NADP-reducing hydrogenase subunit HndC | K18331 | EC:1.12.1.3 | 1 | 1 | 1 | 1 | 1 | 1 | 1 | 1 | 1 | 1 | 1 | 1 | 1 | 1 |
| *hndD* | NADP-reducing hydrogenase subunit HndD | K18332 | EC:1.12.1.3 | 1 | 1 | 1 | 1 | 1 | 1 | 1 | 1 | 1 | 1 | 1 | 1 | 1 | 1 |
| *hycB* | FeS-containing electron transfer protein | K25124 | | 1 | 1 | 1 | 1 | 1 | 1 | 1 | 1 | 1 | 1 | 1 | 1 | 1 | 1 |
| *hydA* | Quinone-reactive Ni/Fe-hydrogenase small subunit | K05927 | EC:1.12.5.1 | 1 | 1 | 1 | 1 | 1 | 1 | 1 | 1 | 1 | 1 | 1 | 1 | 1 | 1 |
| *hydEFG* | NiFe-hydrogenase | | EC 1.12.7.2 | 1 | 1 | 1 | 1 | 1 | 1 | 1 | 1 | 1 | 1 | 1 | 1 | 1 | 1 |
| *hycA* | Formate hydrogenlyase regulatory protein HycA | K15833 | | 0 | 0 | 0 | 0 | 0 | 0 | 0 | 0 | 0 | 0 | 0 | 0 | 0 | 0 |
| *hyaE* | Hydrogenase-1 operon protein HyaE | K03619 | | 1 | 1 | 1 | 1 | 1 | 1 | 1 | 1 | 1 | 1 | 1 | 0 | 0 | 1 |
| *hoxEFU* | Bidirectional [NiFe] hydrogenase diaphorase subunit | K05586, | EC:7.1.1.2 | 1 | 1 | 1 | 1 | 1 | 1 | 1 | 1 | 1 | 1 | 1 | 1 | 1 | 1 |
| *hypABCDEF* | Hydrogenase nickel incorporation protein | K04651, K04652, K046 | | 1 | 1 | 1 | 1 | 1 | 1 | 1 | 1 | 1 | 1 | 1 | 1 | 1 | 1 |
| *echABCDEF* | ech hydrogenase subunits | K14086, K14087, K140 | | 0 | 1 | 0 | 0 | 0 | 0 | 1 | 1 | 0 | 1 | 0 | 1 | 1 | 0 |
| *vhtA* | Methanophenazine hydrogenase, large subunit | K14068 | EC:1.12.98.3 | 0 | 0 | 0 | 0 | 0 | 0 | 0 | 0 | 0 | 0 | 0 | 1 | 1 | 0 |

Supplementary table S3. Annotation details of the genes encoding metabolic pathways of the reconstructed genomes. The genes are sorted by functional groups (in blue) and subgroups (in grey). "0" shows that gene is absent, "1" shows that gene is present.

| | | | | | | | | | | | | | | | | | |
|---|---|---|---|---|---|---|---|---|---|---|---|---|---|---|---|---|---|
| *frhAB* | Coenzyme F420 hydrogenase subunits A and B | K00440 | EC:1.12.98.1 | 1 | 1 | 1 | 1 | 1 | 1 | 1 | 1 | 1 | 1 | 1 | 1 | 1 | 1 |
| *hyaB, hybC* | Hydrogenase large subunit | K06281 | EC:1.12.99.6 | 0 | 0 | 0 | 0 | 0 | 0 | 0 | 0 | 0 | 0 | 0 | 1 | 1 | 0 |
| **Oxidative_phoshorylation** | | | | | | | | | | | | | | | | | |
| *atpE* | F-type H+-transporting ATPase subunit c | K02110 | | 1 | 1 | 1 | 1 | 1 | 1 | 1 | 1 | 1 | 1 | 1 | 0 | 0 | 1 |
| *atpA* | F-type H+/Na+-transporting ATPase subunit alpha | K02111 | EC:7.1.2.2 | 1 | 1 | 1 | 1 | 1 | 1 | 1 | 1 | 1 | 1 | 1 | 0 | 0 | 1 |
| *atpB* | F-type H+/Na+-transporting ATPase subunit beta | K02112 | EC:7.1.2.2 | 1 | 1 | 1 | 1 | 1 | 1 | 1 | 1 | 1 | 1 | 1 | 1 | 1 | 1 |
| *atpH* | F-type H+-transporting ATPase subunit delta | K02113 | | 1 | 1 | 1 | 1 | 1 | 1 | 1 | 1 | 1 | 1 | 1 | 0 | 0 | 0 |
| *atpC* | F-type H+-transporting ATPase subunit epsilon | K02114 | | 1 | 1 | 1 | 1 | 1 | 1 | 1 | 1 | 1 | 1 | 1 | 0 | 0 | 0 |
| *atpG* | F-type H+-transporting ATPase subunit gamma | K02115 | | 1 | 1 | 1 | 1 | 1 | 1 | 1 | 1 | 1 | 1 | 1 | 0 | 0 | 0 |
| *ntpA* | V/A-type H+/Na+-transporting ATPase subunit A | K02117 | EC:7.1.2.2 | 1 | 1 | 1 | 1 | 1 | 1 | 1 | 1 | 1 | 1 | 1 | 1 | 0 | 1 |
| *ntpB* | V/A-type H+/Na+-transporting ATPase subunit B | K02118 | | 1 | 1 | 1 | 1 | 1 | 1 | 1 | 1 | 1 | 1 | 1 | 1 | 0 | 1 |
| *ntpC* | V/A-type H+/Na+-transporting ATPase subunit C | K02119 | | 1 | 1 | 1 | 1 | 1 | 1 | 1 | 1 | 1 | 1 | 1 | 0 | 0 | 0 |
| *ntpD* | V/A-type H+/Na+-transporting ATPase subunit D | K02120 | | 1 | 1 | 1 | 1 | 1 | 1 | 1 | 1 | 1 | 1 | 1 | 1 | 1 | 1 |
| *ntpE* | V/A-type H+/Na+-transporting ATPase subunit E | K02121 | | 1 | 1 | 1 | 1 | 1 | 1 | 1 | 1 | 1 | 1 | 1 | 1 | 1 | 1 |
| *ntpF* | V/A-type H+/Na+-transporting ATPase subunit F | K02122 | | 1 | 1 | 1 | 1 | 1 | 1 | 1 | 1 | 1 | 1 | 1 | 0 | 0 | 0 |
| *nuoE* | NADH-quinone oxidoreductase subunit E | K00334 | EC:7.1.1.2 | 1 | 1 | 1 | 1 | 1 | 1 | 1 | 1 | 1 | 1 | 1 | 1 | 1 | 1 |
| *nuoF* | NADH-quinone oxidoreductase subunit F | K00335 | EC:7.1.1.2 | 1 | 1 | 1 | 1 | 1 | 1 | 1 | 1 | 1 | 1 | 1 | 1 | 1 | 1 |
| *nuoG* | NADH-quinone oxidoreductase subunit G | K00336 | EC:7.1.1.2 | 1 | 1 | 1 | 1 | 1 | 1 | 1 | 1 | 1 | 1 | 1 | 1 | 1 | 1 |
| *ndhA* | NAD(P)H-quinone oxidoreductase subunit 1 | K05572 | EC:7.1.1.2 | 0 | 0 | 0 | 0 | 0 | 0 | 0 | 0 | 0 | 0 | 0 | 1 | 1 | 0 |
| *ndhB* | NAD(P)H-quinone oxidoreductase subunit 2 | K05573 | EC:7.1.1.2 | 1 | 1 | 1 | 1 | 1 | 1 | 1 | 1 | 1 | 1 | 1 | 1 | 1 | 1 |
| *ndhC* | NAD(P)H-quinone oxidoreductase subunit 3 | K05574 | EC:7.1.1.2 | 0 | 0 | 0 | 0 | 0 | 0 | 0 | 0 | 0 | 0 | 0 | 1 | 1 | 0 |
| *ndhD* | NAD(P)H-quinone oxidoreductase subunit 4 | K05575 | EC:7.1.1.2 | 0 | 0 | 0 | 0 | 0 | 0 | 0 | 0 | 0 | 0 | 0 | 1 | 1 | 0 |
| *cydA* | Cytochrome bd ubiquinol oxidase subunit I | K00425 | EC:7.1.1.7 | 0 | 0 | 0 | 0 | 0 | 0 | 0 | 0 | 0 | 0 | 0 | 1 | 1 | 0 |
| *cydB* | Cytochrome bd ubiquinol oxidase subunit II | K00426 | EC:7.1.1.7 | 0 | 0 | 0 | 0 | 0 | 0 | 0 | 0 | 0 | 0 | 0 | 1 | 1 | 0 |
| *SOD2* | Superoxide dismutase, Fe-Mn family | K04564 | EC:1.15.1.1 | 0 | 0 | 0 | 0 | 0 | 0 | 0 | 0 | 0 | 0 | 0 | 1 | 0 | 0 |
| *CCS* | Copper chaperone for superoxide dismutase | K04569 | | 0 | 0 | 0 | 0 | 0 | 0 | 0 | 0 | 0 | 0 | 0 | 0 | 1 | 0 |
| **Nitrogen metabolism** | | | | | | | | | | | | | | | | | |
| **Nitrogen fixation** | | | | | | | | | | | | | | | | | |
| *nifH* | Nitrogenase iron protein NifH | K02588 | EC:1.18.6.1 | 1 | 1 | 1 | 1 | 1 | 1 | 1 | 1 | 0 | 1 | 0 | 0 | 1 | 1 |
| *nifD* | Nitrogenase molybdenum-iron protein alpha chain | K02586 | EC:1.18.6.1 | 1 | 1 | 1 | 1 | 1 | 1 | 1 | 1 | 0 | 1 | 0 | 0 | 1 | 1 |
| *nifK* | Nitrogenase molybdenum-iron protein beta chain | K02591 | EC:1.18.6.1 | 1 | 1 | 1 | 1 | 1 | 1 | 1 | 1 | 0 | 1 | 0 | 0 | 1 | 1 |
| *vnfD* | Vanadium-dependent nitrogenase alpha chain | K22896 | EC:1.18.6.2 | 0 | 1 | 0 | 1 | 1 | 1 | 1 | 1 | 0 | 1 | 0 | 0 | 0 | 1 |
| *vnfK* | Vanadium-dependent nitrogenase beta chain | K22897 | EC:1.18.6.2 | 0 | 1 | 0 | 1 | 1 | 1 | 1 | 1 | 0 | 1 | 0 | 0 | 0 | 1 |
| *vnfH* | Vanadium nitrogenase iron protein | K22899 | | 0 | 1 | 1 | 1 | 1 | 1 | 1 | 1 | 0 | 0 | 0 | 0 | 1 | 1 |
| **Glutamate synthesis** | | | | | | | | | | | | | | | | | |
| *gltBD* | Glutamate synthase (NADPH) large chain | K00265 | EC:1.4.1.13 | 1 | 1 | 1 | 1 | 1 | 1 | 1 | 1 | 1 | 1 | 1 | 1 | 1 | 1 |
| *glnA* | Glutamine synthetase | K01915 | EC:6.3.1.2 | 1 | 1 | 1 | 1 | 1 | 1 | 0 | 1 | 1 | 1 | 1 | 1 | 1 | 1 |
| *gdhA* | Glutamate dehydrogenase (NAD(P)+) | K00261 | EC:1.4.1.3 | 1 | 1 | 1 | 1 | 1 | 1 | 1 | 1 | 1 | 1 | 1 | 0 | 0 | 1 |
| **Urea degradation and ammonium transport** | | | | | | | | | | | | | | | | | |
| *ureC* | Urease subunit alpha | K01428 | EC:3.5.1.5 | 1 | 1 | 1 | 1 | 1 | 1 | 1 | 1 | 1 | 1 | 1 | 0 | 0 | 0 |
| *ureB* | Urease subunit beta | K01429 | EC:3.5.1.5 | 0 | 1 | 0 | 1 | 0 | 0 | 1 | 0 | 0 | 0 | 0 | 0 | 0 | 0 |
| *ureA* | Urease subunit gamma | K01430 | EC:3.5.1.5 | 1 | 1 | 1 | 1 | 1 | 1 | 1 | 1 | 1 | 1 | 1 | 0 | 0 | 0 |
| *urtE* | Urea transport system ATP-binding protein | K11963 | | 1 | 1 | 1 | 1 | 1 | 1 | 1 | 1 | 1 | 1 | 1 | 1 | 1 | 1 |
| *amtB* | Ammonium transporter, Amt family | K03320 | | 1 | 1 | 1 | 1 | 1 | 1 | 1 | 1 | 1 | 1 | 1 | 1 | 1 | 1 |
| **Formate regulation** | | | | | | | | | | | | | | | | | |
| *focA* | Formate transporter | K06212 | | 0 | 1 | 0 | 1 | 1 | 1 | 1 | 1 | 0 | 0 | 0 | 1 | 0 | 1 |
| *pflACE* | Pyruvate formate lyase activating enzyme | K04069 | EC:1.97.1.4 | 1 | 1 | 1 | 1 | 1 | 1 | 1 | 1 | 1 | 1 | 1 | 1 | 1 | 1 |
| *pflX* | Putative pyruvate formate lyase activating enzyme | K04070 | EC:1.97.1.4 | 1 | 1 | 1 | 1 | 1 | 1 | 1 | 1 | 1 | 1 | 1 | 0 | 0 | 1 |
| *fhlA* | Formate hydrogenlyase transcriptional activator | K15836 | | 1 | 1 | 1 | 1 | 1 | 1 | 1 | 1 | 1 | 1 | 1 | 1 | 1 | 1 |
| **Sulfur metabolism** | | | | | | | | | | | | | | | | | |

Supplementary table S3. Annotation details of the genes encoding metabolic pathways of the reconstructed genomes. The genes are sorted by functional groups (in blue) and subgroups (in grey). "0" shows that gene is absent, "1" shows that gene is present.

| | | | | | | | | | | | | | | | | | |
|---|---|---|---|---|---|---|---|---|---|---|---|---|---|---|---|---|---|
| **Sulfate reduction** | | | | | | | | | | | | | | | | | |
| dsrA | Dissimilatory sulfite reductase beta subunit | K11180 | EC:1.8.99.5 | 1 | 1 | 1 | 1 | 1 | 1 | 1 | 1 | 1 | 1 | 1 | 1 | 1 | 1 |
| dsrB | Dissimilatory sulfite reductase alpha subunit | K11181 | EC:1.8.99.5 | 1 | 1 | 1 | 1 | 1 | 1 | 1 | 1 | 1 | 1 | 1 | 1 | 1 | 1 |
| aprA | Adenylylsulfate reductase, subunit A | K00394 | EC:1.8.99.2 | 0 | 0 | 0 | 0 | 0 | 0 | 0 | 0 | 0 | 0 | 0 | 1 | 1 | 1 |
| aprB | Adenylylsulfate reductase, subunit B | K00395 | EC:1.8.99.2 | 0 | 0 | 0 | 0 | 0 | 0 | 0 | 0 | 0 | 0 | 0 | 1 | 1 | 1 |
| asrA | Anaerobic sulfite reductase subunit A | K16950 | | 0 | 0 | 0 | 0 | 0 | 0 | 0 | 0 | 0 | 0 | 0 | 1 | 1 | 1 |
| asrB | Anaerobic sulfite reductase subunit B | K16951 | | 0 | 0 | 0 | 0 | 0 | 0 | 0 | 0 | 0 | 0 | 0 | 1 | 1 | 1 |
| asrC | Anaerobic sulfite reductase subunit C | K00385 | | 0 | 0 | 0 | 0 | 0 | 0 | 0 | 0 | 0 | 0 | 0 | 1 | 1 | 1 |
| sor | Sulfur oxygenase/reductase | K16952 | | 0 | 0 | 0 | 0 | 0 | 0 | 0 | 0 | 0 | 0 | 0 | 0 | 0 | 0 |
| **Cystein biosynthesis** | | | | | | | | | | | | | | | | | |
| cysE | Serine O-acetyltransferase | K00640 | EC:2.3.1.30 | 1 | 1 | 1 | 1 | 1 | 1 | 1 | 1 | 1 | 1 | 1 | 1 | 1 | 1 |
| cysK | Cysteine synthase | K01738 | EC:2.5.1.47 | 1 | 1 | 1 | 1 | 1 | 1 | 1 | 1 | 1 | 1 | 1 | 1 | 1 | 1 |
| cysNC | Bifunctional enzyme CysN/CysC | K00955 | EC:2.7.7.4, 2. | 1 | 1 | 1 | 1 | 1 | 1 | 1 | 1 | 1 | 1 | 1 | 1 | 1 | 1 |
| cysD | Sulfate adenylyltransferase subunit 2 | K00957 | EC:2.7.7.4 | 0 | 0 | 0 | 0 | 0 | 0 | 1 | 0 | 0 | 0 | 0 | 1 | 1 | 1 |
| **Amino acids biosynthesis** | | | | | | | | | | | | | | | | | |
| *hisD* | Histidinol dehydrogenase | K00013 | EC:1.1.1.23 | 1 | 1 | 0 | 1 | 1 | 1 | 1 | 1 | 1 | 1 | 1 | 1 | 0 | 1 |
| *argH* | Argininosuccinate lyase | K01755 | EC:4.3.2.1 | 1 | 1 | 1 | 1 | 1 | 1 | 1 | 1 | 1 | 1 | 1 | 1 | 1 | 1 |
| *lysA* | Diaminopimelate decarboxylase | K01586 | EC:4.1.1.20 | 1 | 1 | 1 | 1 | 1 | 1 | 1 | 1 | 1 | 1 | 1 | 1 | 1 | 1 |
| *serB* | Phosphoserine phosphatase | K01079 | EC:3.1.3.3 | 0 | 0 | 0 | 1 | 1 | 1 | 0 | 0 | 1 | 1 | 1 | 0 | 0 | 0 |
| *thrC* | Threonine synthase | K01733 | EC:4.2.3.1 | 1 | 1 | 1 | 1 | 1 | 1 | 1 | 1 | 1 | 1 | 1 | 0 | 0 | 1 |
| *proC* | Pyrroline-5-carboxylate reductase | K00286 | EC:1.5.1.2 | 1 | 1 | 1 | 1 | 1 | 1 | 1 | 1 | 1 | 1 | 1 | 1 | 1 | 1 |
| *alaA* | Alanine-synthesizing transaminase | K14260 | EC:2.6.1.66 | 0 | 1 | 1 | 1 | 1 | 1 | 1 | 1 | 1 | 1 | 1 | 0 | 1 | 1 |
| *ilvCDE* | Branched-chain amino acid aminotransferase | K00826 | EC:2.6.1.42 | 1 | 1 | 1 | 1 | 1 | 1 | 1 | 1 | 1 | 1 | 1 | 1 | 1 | 1 |
| *leuABC* | 2-isopropylmalate synthase | K01649, | EC:2.3.3.13 | 1 | 1 | 1 | 1 | 1 | 1 | 1 | 1 | 1 | 1 | 1 | 1 | 1 | 1 |
| *metKI* | 5-methyltetrahydropteroyltriglutamate--homocysteine methyltr | K00549 | EC:2.1.1.14 | 1 | 1 | 1 | 1 | 1 | 1 | 1 | 1 | 1 | 1 | 1 | 1 | 1 | 1 |
| *pheA* | Prephenate dehydratase | K04518 | EC:4.2.1.51 | 1 | 1 | 1 | 1 | 1 | 1 | 1 | 1 | 1 | 1 | 1 | 1 | 1 | 1 |
| *tyrAB* | Aromatic-amino-acid transaminase | K00832 | EC:2.6.1.57 | 1 | 1 | 1 | 1 | 1 | 1 | 1 | 1 | 1 | 1 | 1 | 1 | 1 | 1 |
| *trpA* | Tryptophan synthase alpha chain | K01695 | EC:4.2.1.20 | 1 | 1 | 1 | 1 | 1 | 1 | 1 | 1 | 1 | 1 | 1 | 1 | 1 | 1 |
| *aspABC* | Aspartate aminotransferase | K00813 | EC:2.6.1.1 | 1 | 1 | 1 | 1 | 1 | 1 | 1 | 1 | 1 | 1 | 1 | 1 | 1 | 1 |
| *asnB* | Asparagine synthase (glutamine-hydrolysing) | K01953 | EC:6.3.5.4 | 1 | 1 | 1 | 1 | 1 | 1 | 1 | 1 | 1 | 1 | 1 | 1 | 1 | 1 |
| *glyA* | Glycine hydroxymethyltransferase | K00600 | EC:2.1.2.1 | 1 | 1 | 1 | 1 | 1 | 1 | 1 | 1 | 1 | 1 | 1 | 1 | 1 | 1 |
| **Transporters** | | | | | | | | | | | | | | | | | |
| **Peptides (ABC)** | | | | | | | | | | | | | | | | | |
| *ddpD* | Peptide/nickel transport system ATP-binding protein | K02031 | | 0 | 0 | 1 | 1 | 1 | 1 | 0 | 0 | 0 | 0 | 0 | 0 | 0 | 0 |
| *ddpF* | Peptide/nickel transport system ATP-binding protein | K02032 | | 0 | 0 | 1 | 1 | 1 | 1 | 0 | 0 | 0 | 0 | 0 | 0 | 0 | 0 |
| *ABC.PE.P* | Peptide/nickel transport system permease protein | K02033 | | 1 | 1 | 1 | 1 | 1 | 1 | 1 | 1 | 1 | 1 | 1 | 1 | 1 | 1 |
| ABC.PE.P1 | Peptide/nickel transport system permease protein | K02034 | | 1 | 1 | 1 | 1 | 1 | 1 | 1 | 1 | 1 | 1 | 1 | 1 | 1 | 1 |
| ABC.PE.S | Peptide/nickel transport system permease protein | K02035 | | 1 | 1 | 1 | 1 | 1 | 1 | 1 | 1 | 1 | 1 | 1 | 1 | 1 | 1 |
| *oppD* | Oligopeptide transport system ATP-binding protein | K15583 | | 1 | 1 | 0 | 1 | 1 | 1 | 1 | 1 | 0 | 1 | 1 | 1 | 0 | 1 |
| **Transport of inorganic cations/anions** | | | | | | | | | | | | | | | | | |
| *PHO* | Phosphate transporter | K14430 | | 1 | 1 | 1 | 1 | 1 | 1 | 1 | 1 | 1 | 1 | 1 | 1 | 1 | 0 |
| *phoA* | Alkaline phosphatase | K01077 | EC:3.1.3.1 | 0 | 1 | 0 | 1 | 1 | 0 | 0 | 1 | 1 | 1 | 1 | 0 | 0 | 1 |
| *phoU* | Phosphate transport system protein | K02039 | | 1 | 1 | 1 | 1 | 1 | 1 | 1 | 1 | 1 | 1 | 1 | 1 | 1 | 1 |
| *feoB* | Ferrous iron transport protein B | K04759 | | 1 | 1 | 1 | 1 | 1 | 1 | 1 | 1 | 1 | 1 | 1 | 1 | 1 | 1 |
| *cbiMQ* | Cobalt/nickel transport system permease protein | K02008 | | 1 | 1 | 1 | 1 | 0 | 1 | 0 | 0 | 1 | 0 | 1 | 0 | 1 | 0 |
| *corA* | Magnesium transporter | K03284 | | 1 | 0 | 0 | 0 | 0 | 0 | 0 | 0 | 0 | 1 | 1 | 0 | 0 | 0 |
| *copA* | P-type Cu+ transporter | K17686 | | 1 | 1 | 1 | 1 | 1 | 1 | 1 | 1 | 1 | 1 | 1 | 1 | 0 | 1 |
| *thiQ* | Thiamine transport system ATP-binding protein | K02062 | | 1 | 1 | 1 | 1 | 1 | 1 | 1 | 1 | 1 | 1 | 1 | 0 | 0 | 0 |
| *ynjD* | Putative thiamine transport system ATP-binding protein | K05779 | | 1 | 1 | 1 | 1 | 1 | 1 | 1 | 1 | 1 | 1 | 1 | 1 | 1 | 1 |

Supplementary table S3. Annotation details of the genes encoding metabolic pathways of the reconstructed genomes. The genes are sorted by functional groups (in blue) and subgroups (in grey). "0" shows that gene is absent, "1" shows that gene is present.

| | | | | | | | | | | | | | | | | | |
|---|---|---|---|---|---|---|---|---|---|---|---|---|---|---|---|---|---|
| **Substrate transport** | | | | | | | | | | | | | | | | | |
| **PTS: glucose, mannose, fructose, *N*-acetyl-glucosamine** | | | | | | | | | | | | | | | | | |
| *ptsH* | Phosphocarrier protein HPr | K02784 | | 1 | 1 | 1 | 1 | 1 | 1 | 1 | 1 | 1 | 1 | 1 | 1 | 1 | 1 |
| *ptsI* | PTS system enzyme I (PtsI) | K08483 | EC:2.7.3.9 | 1 | 1 | 1 | 1 | 1 | 1 | 1 | 1 | 1 | 1 | 1 | 1 | 1 | 1 |
| *hxlA* | 3-Hexulose-6-phosphate synthase | K08093 | EC:4.1.2.43 | 1 | 1 | 1 | 1 | 1 | 1 | 1 | 1 | 1 | 1 | 1 | 1 | 1 | 1 |
| *manP* | PTS system, mannose-specific EIIBCA component | K25814 | EC:2.7.1.191 | 0 | 0 | 0 | 1 | 1 | 0 | 0 | 0 | 0 | 0 | 0 | 0 | 0 | 0 |
| *GMPP* | Mannose-1-phosphate guanylyltransferase | K00966 | EC:2.7.7.13 | 1 | 1 | 1 | 1 | 1 | 1 | 1 | 1 | 1 | 1 | 1 | 1 | 1 | 1 |
| *fruA* | PTS system, fructose specific EIIB or EIIC component | K02770 | EC:2.7.1.202 | 1 | 1 | 1 | 1 | 1 | 1 | 1 | 1 | 1 | 1 | 1 | 1 | 1 | 1 |
| *fruB* | PTS system, fructose specific EIIA component | K02768 | EC:2.7.1.202 | 1 | 1 | 1 | 1 | 1 | 1 | 1 | 1 | 1 | 1 | 1 | 1 | 1 | 1 |
| **Glucose** | | | | | | | | | | | | | | | | | |
| *glcU* | Glucose uptake protein GlcU | K05340 | | 1 | 1 | 1 | 1 | 1 | 1 | 1 | 1 | 1 | 1 | 1 | 1 | 1 | 1 |
| **Galactose (ABC)** | | | | | | | | | | | | | | | | | |
| *mglC* | Methyl-galactoside transport system permease protein | K10541 | | 1 | 1 | 1 | 1 | 1 | 1 | 1 | 1 | 1 | 1 | 1 | 1 | 1 | 1 |
| *mglA* | Methyl-galactoside transport system ATP-binding protein | K10542 | EC:7.5.2.11 | 1 | 1 | 1 | 1 | 1 | 1 | 1 | 1 | 1 | 1 | 1 | 1 | 1 | 1 |
| *mglB* | Methyl-galactoside transport system substrate-binding protein | K10540 | EC:3.6.3.17 | 1 | 1 | 1 | 1 | 1 | 1 | 1 | 1 | 1 | 1 | 1 | 1 | 1 | 1 |
| **Xylose (ABC)** | | | | | | | | | | | | | | | | | |
| *gguB* | ABC-type xylose transport system, permease component | K10547 | | 1 | 1 | 1 | 1 | 1 | 1 | 1 | 1 | 1 | 1 | 1 | 0 | 0 | 0 |
| *gguA* | ABC-type xylose transport system, ATPase component | K10548 | | 1 | 1 | 1 | 1 | 1 | 1 | 1 | 1 | 1 | 1 | 1 | 1 | 1 | 1 |
| *chvE* | ABC-type xylose transport system, periplasmic component | K10546 | | 1 | 1 | 1 | 1 | 1 | 1 | 1 | 1 | 1 | 1 | 1 | 0 | 0 | 1 |
| *xylF* | ABC-type xylose transport system, periplasmic component | K10543 | | 1 | 1 | 1 | 1 | 1 | 1 | 1 | 1 | 1 | 1 | 1 | 0 | 0 | 1 |
| *xylH* | ABC-type D-xylose transport system permease protein | K10544 | | 1 | 1 | 1 | 1 | 1 | 1 | 1 | 1 | 1 | 1 | 1 | 0 | 0 | 1 |
| *xylG* | ABC-type D-xylose transport system ATP-binding protein | K10545 | EC:7.5.2.10 | 1 | 1 | 1 | 1 | 1 | 1 | 1 | 1 | 1 | 1 | 1 | 1 | 1 | 1 |
| *xylF* | ABC-type D-xylose transport system substrate-binding protein | K10543 | EC:3.6.3.17 | 1 | 1 | 1 | 1 | 1 | 1 | 1 | 1 | 1 | 1 | 1 | 0 | 0 | 1 |
| **Ribose (ABC)** | | | | | | | | | | | | | | | | | |
| *rbsB* | ABC-type ribose transport system substrate-binding protein | K10439 | | 1 | 0 | 1 | 0 | 0 | 0 | 0 | 0 | 1 | 0 | 0 | 0 | 0 | 0 |
| *rbsA* | ABC-type ribose transport system ATP-binding protein | K10441 | EC:7.5.2.7 | 1 | 1 | 1 | 1 | 1 | 1 | 1 | 1 | 1 | 1 | 1 | 1 | 1 | 1 |
| *rbsC* | ABC-type ribose transport system permease protein | K10440 | | 1 | 1 | 1 | 1 | 1 | 1 | 1 | 1 | 1 | 1 | 1 | 0 | 0 | 0 |
| **Arabinose (MFS)** | | | | | | | | | | | | | | | | | |
| *araJ* | MFS transporter, DHA1 family, arabinose polymer utilization pro | K08156 | | 1 | 1 | 1 | 1 | 1 | 1 | 1 | 1 | 1 | 1 | 1 | 1 | 1 | 1 |
| *sotB* | MFS transporter, DHA1 family, L-arabinose/isopropyl-beta-D-thi | K08159 | | 1 | 1 | 0 | 1 | 1 | 1 | 1 | 1 | 0 | 0 | 0 | 1 | 1 | 0 |
| *mdtG, pmrA* | MFS transporter, DHA1 family, multidrug resistance protein | K08161 | | 1 | 1 | 1 | 1 | 1 | 1 | 1 | 1 | 1 | 1 | 1 | 1 | 1 | 1 |
| *mef* | MFS transporter, DHA3 family, macrolide efflux protein | K08217 | | 1 | 1 | 1 | 1 | 1 | 1 | 1 | 1 | 1 | 1 | 1 | 1 | 1 | 1 |
| *MFS* | MFS transporter, SET family, sugar efflux transporter | K03291 | | 1 | 1 | 1 | 1 | 1 | 1 | 1 | 1 | 1 | 1 | 1 | 1 | 1 | 1 |
| **Trehalose (ABC)** | | | | | | | | | | | | | | | | | |
| *treT* | ABC-type trehalose transport system permease protein | K17312 | | 1 | 0 | 1 | 0 | 0 | 0 | 0 | 0 | 1 | 0 | 0 | 0 | 0 | 0 |
| *treU* | ABC-type trehalose transport system permease protein | K17313 | | 1 | 1 | 1 | 1 | 1 | 1 | 1 | 1 | 1 | 1 | 1 | 1 | 1 | 1 |
| *treS* | ABC-type trehalose transport system substrate-binding protein | K17311 | | 1 | 1 | 1 | 1 | 1 | 1 | 1 | 1 | 1 | 1 | 1 | 0 | 0 | 0 |
| *treC* | Trehalose-6-phosphate hydrolase | K01226 | EC:3.2.1.93 | 1 | 0 | 1 | 0 | 0 | 0 | 0 | 0 | 1 | 0 | 0 | 0 | 0 | 0 |
| *treR* | Trehalose operon transcriptional repressor | K03486 | | 1 | 0 | 1 | 0 | 0 | 0 | 0 | 0 | 1 | 0 | 0 | 0 | 0 | 0 |
| **Substrate activation** | | | | | | | | | | | | | | | | | |
| **Hexose** | | | | | | | | | | | | | | | | | |
| *manA* | Mannose-6-phosphate isomerase type 1 | K01809 | EC:5.3.1.8 | 1 | 1 | 1 | 1 | 1 | 1 | 1 | 1 | 1 | 1 | 1 | 0 | 0 | 1 |
| *manC, cpsB* | Mannose-1-phosphate guanylyltransferase RfbM | K00971 | EC:2.7.7.13 | 1 | 1 | 1 | 1 | 1 | 0 | 0 | 0 | 1 | 0 | 0 | 0 | 0 | 1 |
| *scrK* | Fructokinase | K00847 | EC:2.7.1.4 | 1 | 1 | 1 | 1 | 1 | 1 | 1 | 1 | 1 | 1 | 1 | 1 | 0 | 1 |
| *galK* | Galactokinase | K00849 | EC:2.7.1.6 | 1 | 1 | 1 | 1 | 1 | 1 | 1 | 1 | 1 | 1 | 1 | 0 | 0 | 1 |
| *pgmB* | Beta-phosphoglucomutase | K01838 | EC:5.4.2.6 | 1 | 0 | 1 | 0 | 0 | 0 | 0 | 0 | 1 | 0 | 0 | 0 | 0 | 0 |
| *pgm* | Alpha-phosphoglucomutase | K01835 | EC:5.4.2.2 | 1 | 1 | 1 | 1 | 1 | 1 | 1 | 1 | 1 | 1 | 1 | 1 | 1 | 1 |
| ***N*-acetyl-glucosamine** | | | | | | | | | | | | | | | | | |
| *nagA* | N-acetylglucosamine 6-phosphate deacetylase | K01443 | EC:3.5.1.25 | 1 | 1 | 1 | 1 | 1 | 1 | 1 | 1 | 1 | 1 | 1 | 0 | 0 | 1 |
| *nagB* | Glucosamine-6-phosphate deaminase | K02564 | EC:3.5.99.6 | 1 | 1 | 0 | 1 | 1 | 0 | 0 | 0 | 1 | 0 | 0 | 0 | 0 | 0 |

Supplementary table S3. Annotation details of the genes encoding metabolic pathways of the reconstructed genomes. The genes are sorted by functional groups (in blue) and subgroups (in grey). "0" shows that gene is absent, "1" shows that gene is present.

| | | | | | | | | | | | | | | | | | |
|---|---|---|---|---|---|---|---|---|---|---|---|---|---|---|---|---|---|
| **Trehalose** | | | | | | | | | | | | | | | | | |
| *treP* | Alpha,alpha-trehalose phosphorylase | K05342 | EC:2.4.1.64 | 1 | 1 | 1 | 1 | 1 | 1 | 1 | 1 | 1 | 1 | 1 | 0 | 0 | 0 |
| *treC* | Trehalose-6-phosphate hydrolase | K01226 | EC:3.2.1.93 | 1 | 1 | 1 | 1 | 1 | 1 | 1 | 1 | 1 | 1 | 1 | 1 | 1 | 1 |
| *treA, treF* | Alpha,alpha-trehalase | K01194 | EC:3.2.1.28 | 1 | 1 | 1 | 1 | 1 | 1 | 1 | 1 | 1 | 1 | 1 | 1 | 1 | 0 |
| **Pentose** | | | | | | | | | | | | | | | | | |
| *xylA* | D-Xylose isomerase | K01805 | EC:5.3.1.5 | 1 | 1 | 1 | 1 | 1 | 1 | 1 | 1 | 1 | 1 | 1 | 0 | 0 | 0 |
| *xylB* | Xylulokinase | K00854 | EC:2.7.1.17 | 1 | 1 | 1 | 1 | 1 | 1 | 1 | 1 | 1 | 1 | 1 | 0 | 0 | 0 |
| *araA* | L-Arabinose isomerase | K01804 | EC:5.3.1.4 | 1 | 0 | 0 | 0 | 0 | 0 | 0 | 0 | 1 | 0 | 0 | 0 | 0 | 0 |
| *araB* | L-Ribulokinase | K00853 | EC:2.7.1.16 | 1 | 1 | 1 | 1 | 1 | 1 | 1 | 1 | 1 | 1 | 1 | 0 | 0 | 0 |
| *rbsK* | Ribokinase | K00852 | EC:2.7.1.15 | 1 | 1 | 1 | 1 | 1 | 1 | 1 | 1 | 1 | 1 | 1 | 1 | 0 | 1 |
