## Supplemental table S2 for "High-fidelity long-read metagenomics reveals the metabolic functions of dominant termite gut bacteria"

Supplementary table S2. Annotation details of the CAZymes encoding lignocellulose degradation of the reconstructed genomes. In table is shown abundance by a total count number.

| CAZymes | Activities | *Breznakiella ceae strain 1* | *Breznakiella ceae strain 2* | *Breznakiella ceae strain 3* | *Breznakiella ceae strain 4* | *Breznakiella ceae strain 5* | *Breznakiella ceae strain 6* | *Breznakiella ceae strain 7* | *Breznakiella ceae strain 8* | *Breznakiella ceae strain 9* | *Breznakiella ceae strain 10* | *Breznakiella ceae strain 11* | Ca. Fibrimonas strain 1 | Ca. Fibrimonas strain 2 | Chitinispi rillaceae strain 1 |
|---|---|---|---|---|---|---|---|---|---|---|---|---|---|---|---|
| **Cellulose** | | | | | | | | | | | | | | | |
| GH9 | endoglucanase, exo-β-1,4-glucanase, ellobiohydrolase | 0 | 0 | 1 | 0 | 0 | 0 | 1 | 0 | 0 | 0 | 0 | 5 | 4 | 6 |
| GH148 | endo-β-1,4-glucanase | 0 | 0 | 0 | 0 | 0 | 0 | 0 | 0 | 0 | 0 | 0 | 0 | 0 | 1 |
| GH161 | β-1,3-glucan phosphorylase | 0 | 1 | 0 | 0 | 0 | 0 | 0 | 0 | 0 | 0 | 0 | 0 | 0 | 0 |
| **Cellulose and hemicellulose** | | | | | | | | | | | | | | | |
| GH3 | β-glucosidase, xylan 1,4-α-xylosidase | 7 | 3 | 3 | 3 | 3 | 3 | 3 | 4 | 4 | 4 | 5 | 0 | 0 | 0 |
| GH5 | endo-β-1,4-glucanase, endo-β-1,4-xylanase, β-glucosidas | 0 | 0 | 0 | 0 | 0 | 0 | 1 | 0 | 0 | 0 | 0 | 0 | 1 | 0 |
| GH5_2 | chitosanase, endo-β-1,4-glucanase, lichenase / endo-β-1, | 1 | 0 | 0 | 0 | 1 | 1 | 1 | 0 | 0 | 0 | 0 | 4 | 2 | 3 |
| GH5_4 | glucomannan-specific endo-β-1,4-glucanase, endo-β-1,4- | 0 | 1 | 2 | 3 | 1 | 1 | 6 | 2 | 2 | 2 | 2 | 2 | 0 | 0 |
| GH5_10 | xylan β-1,4-xylosidase, endo-β-1,4-mannanase | 0 | 0 | 0 | 0 | 0 | 0 | 0 | 0 | 1 | 0 | 0 | 2 | 0 | 1 |
| GH5_12 | steryl β-glucosidase, β-glucosidase, β-glucosylceramidase | 0 | 1 | 1 | 0 | 0 | 0 | 1 | 1 | 1 | 1 | 1 | 0 | 0 | 0 |
| GH5_25 | endo-β-1,4-glucanase, endo-β-1,4-mannanase, endo-β-1, | 0 | 0 | 0 | 0 | 0 | 0 | 0 | 0 | 1 | 0 | 0 | 0 | 0 | 0 |
| GH5_36 | endo-β-1,4-mannanase | 0 | 0 | 1 | 0 | 0 | 0 | 0 | 0 | 0 | 0 | 0 | 0 | 0 | 0 |
| GH5_37 | endo-β-1,4-glucanase, lichenase / endo-β-1,3-1,4-glucana | 0 | 0 | 0 | 0 | 0 | 0 | 0 | 0 | 0 | 0 | 0 | 1 | 0 | 0 |
| GH5_39 | endo-β-1,4-glucanase | 0 | 0 | 1 | 2 | 2 | 1 | 2 | 2 | 1 | 2 | 2 | 0 | 0 | 0 |
| GH5_40 | endo-β-1,4-mannanase | 0 | 0 | 0 | 0 | 0 | 0 | 0 | 1 | 1 | 0 | 0 | 0 | 0 | 0 |
| GH5_52 | endo-β-1,4-glucanase , exo-β-1,4-glucanase / cellodextrin | 0 | 0 | 0 | 1 | 0 | 0 | 2 | 1 | 1 | 1 | 1 | 1 | 1 | 1 |
| GH8 | chitosanase, endo-β-1,3(4)-glucanase , endo-1,4-β-xylana | 0 | 1 | 0 | 1 | 0 | 1 | 1 | 2 | 0 | 0 | 0 | 2 | 1 | 2 |
| GH16_3 | endo-1,3-β-glucanase, xyloglucanase | 0 | 1 | 0 | 0 | 0 | 0 | 0 | 0 | 0 | 0 | 0 | 0 | 0 | 0 |
| GH30 | β-glucosidase, endo-β-1,4-xylanase | 0 | 3 | 0 | 1 | 1 | 1 | 1 | 1 | 1 | 1 | 0 | 0 | 0 | 0 |
| GH30_1 | β-1,6-glucosidase, β-glucosidase, β-glucosylceramidase | 0 | 1 | 0 | 0 | 0 | 0 | 0 | 0 | 0 | 0 | 0 | 0 | 0 | 0 |
| GH30_8 | Glucuronoarabinoxylan-specific endo-β-1,4-xylanase, end | 0 | 6 | 0 | 1 | 1 | 1 | 1 | 0 | 0 | 0 | 1 | 1 | 0 | 1 |
| GH45 | endoglucanase, endo-xyloglucanase, endo-β-1,4-mannan | 0 | 0 | 0 | 0 | 0 | 0 | 0 | 0 | 0 | 0 | 0 | 2 | 1 | 2 |
| GH51 | endoglucanase, endo-β-1,4-xylanase, β-xylosidase, α-L-ar | 1 | 0 | 0 | 0 | 0 | 0 | 0 | 0 | 0 | 0 | 0 | 1 | 1 | 1 |
| GH112 | D-galactosyl-β-1,4-L-rhamnose phosphorylase | 1 | 0 | 0 | 0 | 0 | 0 | 0 | 0 | 0 | 0 | 0 | 0 | 0 | 0 |
| GH116 | β-glucosidase, β-xylosidase | 0 | 0 | 0 | 0 | 0 | 0 | 0 | 0 | 0 | 0 | 0 | 1 | 2 | 0 |
| **Hemicellulose** | | | | | | | | | | | | | | | |
| CE1 | acetylxylan esterase | 0 | 1 | 1 | 0 | 0 | 1 | 0 | 1 | 0 | 2 | 0 | 0 | 0 | 0 |
| CE2 | acetylxylan esterase, glucuronoyl esterase | 0 | 0 | 0 | 0 | 0 | 0 | 0 | 0 | 0 | 0 | 0 | 1 | 1 | 1 |
| CE3 | acetylxylan esterase | 1 | 0 | 0 | 0 | 0 | 0 | 0 | 0 | 0 | 0 | 0 | 0 | 0 | 0 |
| CE6 | acetylxylan esterase | 0 | 2 | 0 | 1 | 1 | 1 | 1 | 1 | 0 | 1 | 0 | 1 | 1 | 1 |
| CE9 | N-acetylglucosamine-6-phosphate deacetylase | 2 | 1 | 0 | 1 | 0 | 0 | 0 | 0 | 0 | 0 | 0 | 0 | 0 | 0 |
| CE11 | UDP-3-O-acyl-N-acetylglucosamine deacetylase | 0 | 0 | 0 | 0 | 0 | 0 | 0 | 0 | 0 | 0 | 0 | 1 | 1 | 1 |
| CE15 | glucuronoyl esterase | 0 | 0 | 0 | 0 | 0 | 2 | 1 | 1 | 0 | 1 | 1 | 1 | 0 | 1 |
| CE20 | acetylesterase | 4 | 2 | 1 | 2 | 2 | 2 | 4 | 2 | 3 | 3 | 3 | 0 | 0 | 0 |
| GH4 | α-glucosidase, α-galactosidase | 2 | 0 | 0 | 1 | 2 | 1 | 1 | 2 | 0 | 2 | 2 | 0 | 0 | 0 |
| GH10 | endo-1,4-β-xylanase | 2 | 4 | 3 | 7 | 5 | 9 | 8 | 7 | 3 | 5 | 6 | 0 | 0 | 0 |
| GH11 | endo-β-1,4-xylanase, exo-1,4-β-xylosidase | 1 | 2 | 1 | 6 | 3 | 3 | 2 | 5 | 2 | 4 | 2 | 2 | 0 | 3 |
| GH13 | β-glucosidase | 2 | 2 | 0 | 1 | 1 | 1 | 2 | 1 | 1 | 1 | 2 | 0 | 0 | 0 |
| GH13_3 | α-1,4-glucan: phosphate α-maltosyltransferase | 0 | 0 | 0 | 0 | 0 | 0 | 0 | 0 | 0 | 0 | 0 | 1 | 0 | 1 |
| GH13_9 | α-1,4-glucan branching enzyme | 0 | 0 | 0 | 0 | 0 | 0 | 0 | 0 | 0 | 0 | 0 | 1 | 0 | 1 |

Supplementary table S2. Annotation details of the CAZymes encoding lignocellulose degradation of the reconstructed genomes. In table is shown abundance by a total count number.

| | | | | | | | | | | | | | | | |
|---|---|---|---|---|---|---|---|---|---|---|---|---|---|---|---|
| GH13_11 | α-glucosidase | 1 | 1 | 1 | 1 | 1 | 1 | 1 | 1 | 1 | 1 | 1 | 0 | 0 | 0 |
| GH13_14 | α-glucosidase, α-amylase | 0 | 1 | 0 | 0 | 0 | 0 | 0 | 0 | 0 | 0 | 0 | 0 | 0 | 0 |
| GH13_20 | α-amylase, cyclic α-1,6-maltosyl-maltose hydrolase | 0 | 0 | 1 | 1 | 0 | 0 | 1 | 0 | 0 | 0 | 1 | 1 | 1 | 1 |
| GH13_23 | α-glucosyltransferase, α-glucosidase | 0 | 0 | 0 | 0 | 0 | 0 | 0 | 0 | 0 | 0 | 0 | 1 | 1 | 0 |
| GH13_31 | α-glucosidase, α-amylase, α-transglucosidase / α-glucosyl | 1 | 0 | 2 | 0 | 1 | 1 | 1 | 2 | 1 | 2 | 1 | 0 | 0 | 0 |
| GH13_36 | α-amylase | 1 | 2 | 0 | 0 | 0 | 0 | 0 | 0 | 0 | 0 | 0 | 0 | 0 | 0 |
| GH26 | endo-β-1,3-1,4-glucanase, β-1,3-xylanase, β-mannanase | 0 | 0 | 0 | 0 | 0 | 0 | 2 | 0 | 1 | 0 | 0 | 0 | 0 | 1 |
| GH27 | α-galactosidase, α-L-arabinopyranosidase | 0 | 0 | 0 | 0 | 0 | 0 | 0 | 0 | 1 | 0 | 0 | 0 | 0 | 0 |
| GH35 | β-galactosidase | 1 | 0 | 0 | 0 | 0 | 0 | 0 | 0 | 0 | 0 | 0 | 0 | 0 | 0 |
| GH39 | β-xylosidase, α-L-arabinofuranosidase | 1 | 1 | 1 | 1 | 1 | 1 | 1 | 0 | 1 | 0 | 1 | 0 | 0 | 0 |
| GH43_1 | β-xylosidase, α-L-arabinofuranosidase, xylanase | 0 | 1 | 1 | 2 | 1 | 1 | 1 | 1 | 1 | 2 | 1 | 0 | 0 | 0 |
| GH43_2 | α-L-arabinofuranosidase, xylan β-1,4-xylosidase | 0 | 0 | 0 | 0 | 0 | 1 | 1 | 1 | 1 | 0 | 0 | 0 | 0 | 0 |
| GH43_10 | xylan β-1,4-xylosidase, α-L-arabinofuranosidase | 1 | 1 | 0 | 0 | 0 | 0 | 0 | 0 | 0 | 0 | 0 | 0 | 0 | 0 |
| GH43_12 | xylan β-1,4-xylosidase, α-L-arabinofuranosidase, β-1,3-xyl | 0 | 0 | 1 | 1 | 1 | 1 | 1 | 1 | 1 | 1 | 1 | 0 | 0 | 0 |
| GH43_16 | xylan β-1,4-xylosidase, α-L-arabinofuranosidase, endo-β-1 | 0 | 3 | 0 | 0 | 0 | 0 | 0 | 0 | 0 | 0 | 0 | 1 | 0 | 0 |
| GH43_17 | α-L-arabinofuranosidase, endo-β-1,4-xylanase | 0 | 0 | 0 | 0 | 0 | 0 | 0 | 0 | 0 | 0 | 0 | 0 | 1 | 0 |
| GH43_35 | xylan β-1,4-xylosidase, α-L-arabinofuranosidase, endo-β-1 | 1 | 0 | 0 | 1 | 1 | 0 | 0 | 0 | 0 | 1 | 0 | 0 | 0 | 0 |
| GH52 | β-xylosidase | 0 | 0 | 0 | 0 | 0 | 0 | 0 | 0 | 0 | 0 | 0 | 0 | 0 | 1 |
| GH55 | exo-β-1,3-glucanase, endo-β-1,3-glucanase | 0 | 1 | 0 | 0 | 0 | 0 | 0 | 0 | 0 | 0 | 0 | 0 | 0 | 0 |
| GH67 | α-glucuronidase | 4 | 1 | 0 | 0 | 0 | 0 | 1 | 1 | 1 | 1 | 0 | 0 | 0 | 0 |
| GH74 | endoglucanase, xyloglucanase | 0 | 0 | 0 | 0 | 0 | 0 | 1 | 0 | 0 | 0 | 0 | 0 | 0 | 1 |
| GH115 | xylan α-1,2-glucuronidase | 0 | 2 | 0 | 1 | 1 | 1 | 0 | 0 | 0 | 0 | 1 | 0 | 0 | 0 |
| GH120 | β-xylosidase | 1 | 0 | 0 | 0 | 0 | 0 | 0 | 0 | 0 | 1 | 1 | 0 | 0 | 0 |
| GH127 | β-L-arabinofuranosidase | 1 | 0 | 0 | 0 | 0 | 0 | 0 | 0 | 0 | 0 | 0 | 0 | 0 | 0 |
| GH128 | β-1,3-glucanase | 0 | 3 | 0 | 0 | 0 | 0 | 0 | 0 | 0 | 0 | 0 | 0 | 0 | 0 |
| GH130 | β-1,4-mannosylglucose phosphorylase | 1 | 0 | 1 | 1 | 1 | 1 | 2 | 1 | 1 | 1 | 1 | 0 | 0 | 0 |
| GH141 | α-L-fucosidase, endo-β-1,4-xylanase | 0 | 0 | 0 | 0 | 0 | 0 | 0 | 0 | 0 | 1 | 0 | 0 | 0 | 0 |
| GH142 | β-L-arabinofuranosidase | 1 | 1 | 1 | 1 | 1 | 1 | 1 | 1 | 0 | 1 | 1 | 0 | 0 | 0 |
| GH165 | β-galactosidase | 2 | 0 | 0 | 0 | 0 | 0 | 0 | 0 | 0 | 0 | 0 | 0 | 0 | 0 |
| **Lignin** | | | | | | | | | | | | | | | |
| AA4 | vanillyl-alcohol oxidase (EC 1.1.3.38) | 1 | 0 | 0 | 0 | 0 | 0 | 0 | 0 | 0 | 0 | 0 | 0 | 0 | 0 |
| EC:1.1.99. | cellobiose dehydrogenase | 7 | 2 | 2 | 6 | 4 | 3 | 3 | 3 | 4 | 2 | 4 | 4 | 6 | 6 |
| EC:1.1.3.1 | pyranose oxidase | 2 | 0 | 1 | 0 | 0 | 0 | 1 | 1 | 1 | 1 | 1 | 2 | 2 | 2 |
| EC:1.10.3. | laccase | 0 | 0 | 1 | 0 | 0 | 0 | 0 | 0 | 0 | 1 | 1 | 0 | 0 | 0 |
| EC:1.11.1. | manganese peroxidase | 1 | 0 | 0 | 0 | 1 | 0 | 0 | 0 | 0 | 0 | 0 | 0 | 0 | 0 |
| EC:3.1.1.1 | (4-O-methyl)-D-glucuronate---lignin esterase | 0 | 0 | 0 | 0 | 0 | 2 | 1 | 1 | 0 | 1 | 1 | 1 | 0 | 1 |
| **Chitin** | | | | | | | | | | | | | | | |
| CE4 | acetyl xylan esterase, chitin deacetylase | 2 | 1 | 1 | 3 | 1 | 2 | 1 | 2 | 2 | 2 | 1 | 0 | 1 | 2 |
| GH18 | chitinase | 1 | 2 | 1 | 1 | 1 | 1 | 1 | 1 | 1 | 1 | 1 | 1 | 1 | 0 |
| GH20 | β-hexosaminidase | 1 | 1 | 0 | 1 | 0 | 0 | 1 | 0 | 0 | 0 | 0 | 0 | 0 | 1 |
| GH23 | lysozyme type G, peptidoglycan lyase, chitinase | 1 | 3 | 2 | 3 | 3 | 4 | 3 | 3 | 3 | 3 | 3 | 4 | 4 | 6 |
| **Oligosaccharides** | | | | | | | | | | | | | | | |
| GH1 | exo-β-1,4glucanase, β-glucosidase, β-galactosidase, β- ma | 1 | 0 | 1 | 1 | 1 | 1 | 0 | 0 | 0 | 0 | 0 | 0 | 0 | 1 |
| GH2 | β-galactosidases, β-glucuronidases, β-mannosidase | 5 | 1 | 0 | 1 | 1 | 1 | 1 | 1 | 1 | 1 | 0 | 1 | 1 | 0 |

Supplementary table S2. Annotation details of the CAZymes encoding lignocellulose degradation of the reconstructed genomes. In table is shown abundance by a total count number.

| | | | | | | | | | | | | | | | |
|---|---|---|---|---|---|---|---|---|---|---|---|---|---|---|---|
| GH31 | α-glucosidase, α-xylosidase, α-galactosidase, α-mannosid | 1 | 0 | 0 | 0 | 0 | 0 | 1 | 0 | 1 | 0 | 0 | 0 | 0 | 0 |
| GH42 | β-galactosidase, α-L-arabinopyranosidase | 1 | 1 | 0 | 0 | 0 | 0 | 1 | 0 | 0 | 0 | 0 | 0 | 0 | 0 |
| GH57 | α-galactosidase, α-amylase | 2 | 2 | 2 | 2 | 2 | 2 | 2 | 2 | 2 | 2 | 2 | 1 | 1 | 5 |
| GH94 | cellobiose/chitobiose phosphorylase | 2 | 1 | 2 | 2 | 2 | 4 | 2 | 2 | 2 | 2 | 2 | 1 | 0 | 2 |
| GH95 | α-L-fucosidase, α-L-galactosidase | 0 | 0 | 0 | 0 | 0 | 0 | 1 | 0 | 0 | 0 | 0 | 0 | 0 | 0 |
| GH105 | unsaturated rhamnogalacturonyl hydrolase | 3 | 0 | 0 | 0 | 0 | 0 | 0 | 0 | 0 | 0 | 0 | 0 | 0 | 0 |
| GH109 | α-N-acetylgalactosaminidase | 2 | 0 | 0 | 2 | 1 | 0 | 3 | 3 | 1 | 3 | 1 | 0 | 0 | 0 |
| **Pectinases** | | | | | | | | | | | | | | | |
| GH28 | polygalacturonase | 1 | 0 | 0 | 0 | 0 | 0 | 0 | 0 | 0 | 0 | 0 | 0 | 0 | 0 |
| GH53 | endo-β-1,4-galactanase | 0 | 0 | 0 | 0 | 0 | 0 | 0 | 0 | 0 | 0 | 0 | 0 | 1 | 1 |
| GH63 | α-glucosidase | 1 | 0 | 0 | 0 | 0 | 0 | 0 | 0 | 0 | 0 | 0 | 0 | 0 | 0 |
| GH106 | α-L-rhamnosidase | 1 | 0 | 0 | 0 | 0 | 0 | 0 | 0 | 0 | 0 | 0 | 0 | 0 | 1 |
| PL1 | pectin lyase | 0 | 0 | 0 | 0 | 0 | 0 | 0 | 0 | 0 | 0 | 0 | 1 | 0 | 0 |
| PL11 | rhamnogalacturonan endo- and exolyase | 1 | 0 | 0 | 0 | 0 | 0 | 0 | 0 | 0 | 0 | 0 | 1 | 0 | 0 |
| PL11_1 | poly(β-mannuronate) lyase, β-1,4-glucuronan lyase | 0 | 0 | 0 | 0 | 0 | 0 | 0 | 0 | 0 | 0 | 0 | 0 | 1 | 1 |
| PL14_3 | poly(β-mannuronate) lyase, β-1,4-glucuronan lyase | 0 | 0 | 0 | 0 | 0 | 0 | 0 | 0 | 0 | 0 | 0 | 0 | 1 | 0 |
| **Others** | | | | | | | | | | | | | | | |
| GH33 | sialidase or neuraminidase | 1 | 0 | 0 | 0 | 0 | 0 | 0 | 0 | 0 | 0 | 0 | 0 | 0 | 0 |
| GH73 | peptidoglycan hydrolase, lysozyme | 1 | 1 | 1 | 1 | 1 | 1 | 1 | 1 | 1 | 1 | 1 | 0 | 0 | 0 |
| GH77 | amylomaltase | 1 | 1 | 1 | 1 | 1 | 1 | 1 | 1 | 1 | 1 | 1 | 1 | 1 | 1 |
| GH88 | d-4,5-unsaturated β-glucuronyl hydrolase | 1 | 0 | 0 | 0 | 0 | 0 | 0 | 0 | 0 | 0 | 0 | 0 | 0 | 0 |
| GH154 | β-glucuronidase | 2 | 0 | 0 | 0 | 0 | 0 | 0 | 0 | 0 | 0 | 0 | 0 | 0 | 0 |
| GH171 | peptidoglycan β-N-acetylmuramidase | 1 | 0 | 0 | 0 | 0 | 0 | 0 | 0 | 0 | 0 | 0 | 1 | 0 | 1 |
| **Carbohydrate-binding module** | | | | | | | | | | | | | | | |
| CBM4 | xylan, β-1,3-glucan, β-1,3-1,4-glucan, β-1,6-glucan and an | 0 | 0 | 0 | 0 | 0 | 2 | 0 | 0 | 0 | 0 | 0 | 2 | 1 | 6 |
| CBM6 | Amorphous cellulose and β-1,4-xylan-binding | 0 | 1 | 0 | 0 | 0 | 0 | 0 | 0 | 0 | 0 | 0 | 1 | 0 | 1 |
| CBM9 | Xylan-binding | 0 | 1 | 0 | 0 | 0 | 0 | 1 | 0 | 0 | 0 | 1 | 0 | 0 | 1 |
| CBM11 | Cellulose and chitin-binding | 0 | 0 | 0 | 0 | 0 | 0 | 0 | 0 | 0 | 0 | 0 | 2 | 2 | 1 |
| CBM13 | Xylan-binding | 0 | 2 | 0 | 0 | 0 | 0 | 0 | 0 | 0 | 0 | 1 | 0 | 0 | 1 |
| CBM27 | Mannan-binding | 0 | 0 | 0 | 0 | 0 | 0 | 1 | 0 | 0 | 0 | 0 | 0 | 0 | 0 |
| CBM30 | Cellulose-binding | 0 | 0 | 0 | 0 | 0 | 0 | 0 | 0 | 0 | 0 | 0 | 2 | 2 | 1 |
| CBM32 | Galactose-binding | 0 | 0 | 0 | 0 | 0 | 0 | 0 | 0 | 0 | 0 | 0 | 1 | 0 | 1 |
| CBM35 | Xylan, mannan and β-galactan-binding | 0 | 0 | 0 | 0 | 0 | 0 | 0 | 0 | 0 | 0 | 0 | 0 | 0 | 1 |
| CBM36 | Xylan-binding | 0 | 2 | 0 | 2 | 0 | 0 | 0 | 2 | 0 | 0 | 0 | 0 | 0 | 0 |
| CBM46 | Cellulose-binding | 0 | 2 | 0 | 1 | 0 | 0 | 1 | 3 | 0 | 0 | 0 | 0 | 0 | 0 |
| CBM48 | Glycogen-binding | 0 | 0 | 0 | 0 | 1 | 1 | 1 | 1 | 0 | 0 | 0 | 1 | 0 | 1 |
| CBM51 | Galactose-binding | 0 | 0 | 0 | 0 | 0 | 0 | 0 | 0 | 0 | 0 | 0 | 2 | 1 | 0 |
| CBM91 | Xylan-binding | 1 | 1 | 1 | 1 | 1 | 0 | 1 | 1 | 1 | 1 | 1 | 0 | 0 | 0 |
| **Transferase** | | | | | | | | | | | | | | | |
| GT1 | Various | 0 | 0 | 0 | 0 | 0 | 0 | 0 | 1 | 0 | 0 | 0 | 0 | 0 | 0 |
| GT2 | cellulose synthase, chitin synthase, β-1,3-glucan synthase | 3 | 9 | 3 | 4 | 4 | 3 | 8 | 8 | 7 | 3 | 5 | 20 | 15 | 15 |
| GT4 | N-acetylglucosamine (GlcNAc) | 5 | 3 | 5 | 8 | 6 | 6 | 9 | 6 | 7 | 5 | 6 | 8 | 10 | 8 |
| GT5 | glycogen and starch glucosyltransferase | 1 | 2 | 1 | 1 | 1 | 2 | 1 | 1 | 1 | 1 | 1 | 2 | 1 | 2 |
| GT8 | lipopolysaccharide α-1,3-galactosyltransferase | 0 | 0 | 1 | 0 | 0 | 0 | 0 | 0 | 0 | 0 | 0 | 1 | 1 | 0 |

Supplementary table S2. Annotation details of the CAZymes encoding lignocellulose degradation of the reconstructed genomes. In table is shown abundance by a total count number.

| | | | | | | | | | | | | | | | |
|---|---|---|---|---|---|---|---|---|---|---|---|---|---|---|---|
| GT10 | galactoside α-1,3/1,4-L-fucosyltransferase | 0 | 0 | 0 | 0 | 1 | 0 | 0 | 0 | 0 | 0 | 0 | 0 | 0 | 0 |
| GT11 | galactoside α-1,2-L-fucosyltransferase | 0 | 0 | 0 | 0 | 0 | 0 | 0 | 0 | 0 | 0 | 0 | 0 | 0 | 1 |
| GT19 | lipid-A-disaccharide synthase | 0 | 0 | 0 | 0 | 0 | 0 | 0 | 0 | 0 | 0 | 0 | 1 | 0 | 1 |
| GT21 | ceramide β-glucosyltransferase | 1 | 0 | 0 | 0 | 0 | 0 | 0 | 0 | 0 | 0 | 0 | 0 | 0 | 1 |
| GT26 | β-1,4-glucosyltransferase, β-1,4-galactosyltransferase, β-l | 1 | 1 | 1 | 1 | 1 | 1 | 1 | 1 | 1 | 1 | 1 | 0 | 0 | 0 |
| GT27 | polypeptide α-N-acetylgalactosaminyltransferase | 0 | 0 | 0 | 0 | 0 | 0 | 1 | 0 | 0 | 0 | 0 | 0 | 0 | 0 |
| GT28 | 1,2-diacylglycerol 3-β-galactosyltransferase, 1,2-diacylgly | 2 | 1 | 1 | 1 | 1 | 1 | 1 | 1 | 1 | 1 | 1 | 1 | 1 | 1 |
| GT32 | α-1,6-mannosyltransferase, α-1,4-N-acetylglucosaminyltr | 0 | 0 | 1 | 1 | 0 | 0 | 1 | 0 | 1 | 0 | 2 | 1 | 2 | 2 |
| GT35 | glycogen or starch phosphorylase | 1 | 1 | 1 | 1 | 1 | 1 | 1 | 1 | 1 | 1 | 1 | 1 | 1 | 2 |
| GT51 | murein polymerase | 2 | 2 | 2 | 3 | 2 | 2 | 2 | 2 | 2 | 2 | 2 | 4 | 3 | 4 |
| GT92 | N-glycan core α-1,6-fucoside β-1,4-galactosyltransferase | 0 | 0 | 2 | 0 | 0 | 0 | 0 | 0 | 0 | 0 | 0 | 0 | 0 | 0 |
| GT104 | arginine α-L-rhamnosyltransferase | 0 | 0 | 0 | 0 | 0 | 0 | 0 | 0 | 0 | 0 | 0 | 0 | 0 | 2 |
| GT111 | β-1,3-galactofuranosyltransferase | 0 | 0 | 0 | 0 | 0 | 0 | 0 | 0 | 0 | 0 | 0 | 1 | 1 | 0 |
