## Supplemental table S4 for "High-fidelity long-read metagenomics reveals the metabolic functions of dominant termite gut bacteria"

Supplementary table S4. Comparison of protein sequence identity and horizontal gene transfer events between the reconstracted genomes.

| qseqid | sseqid | pident | length | evalue | qcovhsp | KO | KO definition |
|---|---|---|---|---|---|---|---|
| Chitinispirillaceae_strain_1_141 | Breznakiellaceae_strain_1_31 | 93.6 | 204 | 1.04E-138 | 100 | no annotation | |
| Chitinispirillaceae_strain_1_1182 | Breznakiellaceae_strain_6_2370 | 90.3 | 145 | 2.43E-99 | 100 | K00196 | anaerobic carbon-monoxide dehydrogenase iron sulfur subunit |
| Chitinispirillaceae_strain_1_1182 | Breznakiellaceae_strain_5_65 | 90.3 | 145 | 2.43E-99 | 100 | K00197 | anaerobic carbon-monoxide dehydrogenase iron sulfur subunit |
| Chitinispirillaceae_strain_1_1486 | Breznakiellaceae_strain_5_1066 | 91.1 | 270 | 1.74E-178 | 98.9 | K02588 | nitrogenase iron protein NifH |
| Chitinispirillaceae_strain_1_1988 | Breznakiellaceae_strain_2_2555 | 91.7 | 503 | 0 | 100 | K03427 | type I restriction enzyme M protein [EC:2.1.1.72] |
| Chitinispirillaceae_strain_1_2917 | Breznakiellaceae_strain_6_1408 | 91.4 | 267 | 1.90E-179 | 100 | no annotation | |
| Chitinispirillaceae_strain_1_3124 | Breznakiellaceae_strain_2_2330 | 91.9 | 149 | 1.97E-98 | 100 | no annotation | |
| Chitinispirillaceae_strain_1_1486 | Ca_Fibrimonas_strain_2_253 | 91.9 | 270 | 2.65E-181 | 98.9 | K02588 | nitrogenase iron protein NifH |
| Chitinispirillaceae_strain_1_1825 | Ca_Fibrimonas_strain_1_682 | 93.5 | 291 | 1.74E-195 | 100 | | |
| Chitinispirillaceae_strain_1_1826 | Ca_Fibrimonas_strain_1_683 | 94.3 | 279 | 9.66E-187 | 100 | K02050 | NitT/TauT family transport system permease protein |
| Chitinispirillaceae_strain_1_2917 | Ca_Fibrimonas_strain_2_1988 | 93.3 | 267 | 4.11E-182 | 100 | no annotation | |
| Ca_Fibrimonas_strain_2_253 | Breznakiellaceae_strain_5_1066 | 97.4 | 273 | 4.75E-192 | 100 | K02588 | nitrogenase iron protein NifH |
| Ca_Fibrimonas_strain_2_1496 | Breznakiellaceae_strain_11_114 | 90.8 | 349 | 6.81E-234 | 99.4 | no annotation | |
| Ca_Fibrimonas_strain_2_1638 | Breznakiellaceae_strain_4_1459 | 94.2 | 172 | 7.38E-121 | 100 | no annotation | |
| Ca_Fibrimonas_strain_2_1648 | Breznakiellaceae_strain_11_128 | 92.2 | 231 | 1.52E-158 | 100 | no annotation | |
| Ca_Fibrimonas_strain_2_1716 | Breznakiellaceae_strain_4_644 | 93.9 | 164 | 5.18E-115 | 99.4 | no annotation | |
| Ca_Fibrimonas_strain_2_1988 | Breznakiellaceae_strain_6_1408 | 94.4 | 267 | 1.85E-186 | 100 | no annotation | |
| Ca_Fibrimonas_strain_2_2141 | Breznakiellaceae_strain_7_1460 | 93.4 | 437 | 2.57E-302 | 100 | no annotation | |
| Ca_Fibrimonas_strain_2_2318 | Breznakiellaceae_strain_6_1588 | 93.8 | 437 | 3.60E-313 | 100 | K01845 | glutamate-1-semialdehyde 2,1-aminomutase [EC:5.4.3.8] |
| Ca_Fibrimonas_strain_1_2433 | Breznakiellaceae_strain_11_128 | 92.2 | 231 | 2.16E-158 | 100 | no annotation | |
| Ca_Fibrimonas_strain_1_2434 | Breznakiellaceae_strain_1_4129 | 92.3 | 439 | 3.38E-287 | 100 | no annotation | |
